# Differential reliance of CTD-nuclear envelope phosphatase 1 on its regulatory subunit in ER lipid synthesis and storage

**DOI:** 10.1101/2023.10.12.562096

**Authors:** Jake W. Carrasquillo Rodríguez, Onyedikachi Uche, Shujuan Gao, Shoken Lee, Michael V. Airola, Shirin Bahmanyar

**Affiliations:** Department of Molecular, Cellular and Developmental Biology, Yale University, New Haven, CT 06511; Department of Biochemistry and Cell Biology, Stony Brook University, Stony Brook NY 11794

## Abstract

The endoplasmic reticulum (ER) is the site for the synthesis of the major membrane and storage lipids. Lipin 1 produces diacylglycerol, the lipid intermediate critical for the synthesis of both membrane and storage lipids in the ER. CTD-Nuclear Envelope Phosphatase 1 (CTDNEP1) regulates lipin 1 to restrict ER membrane synthesis, but its role in lipid storage in mammalian cells is unknown. Here, we show that the ubiquitin-proteasome degradation pathway controls the levels of ER/nuclear envelope-associated CTDNEP1 to regulate ER membrane synthesis through lipin 1. The N-terminus of CTDNEP1 is an amphipathic helix that targets to the ER, nuclear envelope and lipid droplets. We identify key residues at the binding interface of CTDNEP1 with its regulatory subunit NEP1R1 and show that they facilitate complex formation *in vivo* and *in vitro*. We demonstrate a role for NEP1R1 in temporarily shielding CTDNEP1 from proteasomal degradation to regulate lipin 1 and restrict ER size. Unexpectedly, we found that NEP1R1 is not required for CTDNEP1’s role in restricting lipid droplet biogenesis. Thus, the reliance of CTDNEP1 function on its regulatory subunit differs during ER membrane synthesis and lipid storage. Together, our work provides a framework into understanding how the ER regulates lipid synthesis and storage under fluctuating conditions.

## Introduction

Membrane bound organelles have characteristic shapes and compositions (Voeltz and Prinz, 2007). Fluctuations in nutrient availability, cell cycle stage, and stress conditions can dramatically change the compositions of membrane bound organelles (Jackowski, 1994; Efeyan et al., 2015). Transcriptional and post-translational mechanisms sense and respond to changing conditions to restore organelle homeostasis and protect against potential toxic effects (Stevenson et al., 2016). Alterations in these mechanisms disrupt cellular homeostasis, which is a hallmark of many pathological disorders, including cancer (Menendez and Lupu, 2007; Currie et al., 2013).

The endoplasmic reticulum (ER) is the largest organelle in cells that carries out many essential cellular functions (Baumann and Walz, 2001). The ER is made up of tubules, sheets and the nuclear envelope (NE) that enclose a single continuous lumen (Friedman et al., 2011). The ER is the site of synthesis of membrane embedded and soluble proteins destined for secretion, and much of the protein machinery in the ER functions in the synthesis, processing and trafficking of secreted proteins. Protein quality control mechanisms recognize misfolded proteins and target them for degradation by the ER-associated degradation pathway (Mehrtash and Hochstrasser, 2018; Christianson et al., 2023). The AAA^+^-ATPase p97 cooperates with the proteasome to extract membrane proteins for their subsequent degradation (Meyer et al., 2012).

The ER is also the site of *de novo* synthesis of most membrane and storage lipids (Fagone and Jackowski, 2009). Enzymes embedded or peripherally associated with the surface of the ER carry out a series of sequential reactions to produce the major membrane phospholipids (e.g. phosphatidylcholine and phosphatidylethanolamine) from diacylglycerol (DAG) and a more minor pool of lipid (e.g. phosphatidylinositol) from phosphatidic acid (PA). Activated fatty acids combine with lysophosphatidic acid (LPA) to produce PA, which is dephosphorylated to generate DAG. DAG can also be converted to triglycerides (TAG) (Walther and Farese, 2012). TAGs accumulate in specialized storage organelles called lipid droplets (LDs), which are derived from the ER and protect cells from buildup of excess fatty acids that can negatively effect membrane function (Coleman et al., 2000; Olzmann and Carvalho, 2019). Feedback mechanisms maintain ER lipid homeostasis by sensing and responding to biochemical fluxes in lipid levels (Goldstein et al., 2006; Olzmann and Carvalho, 2019). These feedback mechanisms control the quantity of lipid enzymes through transcriptional networks and protein degradation machinery (Stevenson et al., 2016).

As the major supplier of membrane lipids for organelles, the ER must maintain its overall structure despite constant flow of lipids away from the ER. How the production of membranes in the ER is regulated to control its size and dynamics in animal cells is not fully understood. Furthermore, the upstream mechanisms that control the switch between lipid storage and membrane production in the ER are not well understood, particularly in metazoans that use the same lipid intermediate (DAG) for both reactions (Coleman and Lee, 2004).

Here, we focused on the protein phosphatase CTD-nuclear envelope phosphatase 1 (CTDNEP1, formerly known as Dullard; Nem1 in *S. cerevisiae*, CNEP-1 in *C. elegans*) because of its conserved role in controlling the production of ER and nuclear membranes (Siniossoglou et al., 1998; Santos-Rosa et al., 2005; O’Hara et al., 2006; Kim et al., 2007; Han et al., 2012; Bahmanyar et al., 2014; Jacquemyn et al., 2021; Merta et al., 2021). CTDNEP1 is enriched at the nuclear envelope and maintains a stable, dephosphorylated pool of the phosphatidic acid phosphatase lipin 1, the main enzyme that produces diacylglycerol (DAG) in the ER (Siniossoglou et al., 1998; Han et al., 2012; Merta et al., 2021; Lee et al., 2023). CTDNEP1 opposes stimulatory signals from nutrient sensing pathways that retain phosphorylated lipin 1 in the cytoplasm (Merta et al., 2021). Dephosphorylated lipin 1 is nuclear-localized and has a role in restricting the transcription of fatty acid synthesis genes (Peterson et al., 2011; Merta et al., 2021). In CTDNEP1 knockout cells, lipin 1 is hyperphosphorylated and there is an increased biochemical flux of fatty acid synthesis that then results in excess ER membrane production (Merta et al., 2021). It has been shown that dephosphorylated lipin 1 has increased association with membranes (Harris et al., 2007; Eaton et al., 2013); however, DAG levels are unchanged in CTDNEP1 knockout cells (Merta et al., 2021), which may be because of increasing fatty acid availability also promotes lipin 1’s membrane association to turn over PA (Harris et al., 2007; Eaton et al., 2013;). Thus, CTDNEP1 control of lipin 1 is coordinated with the biochemical flux of *de novo* fatty acid synthesis to match DAG synthesis. How CTDNEP1 is regulated to accommodate changing demands for ER lipid synthesis in mammalian cells is not known.

Important work in budding yeast has elucidated mechanisms that regulate the conserved lipin activation pathway to control ER lipid homeostasis (Santos-Rosa et al., 2005; O’Hara et al., 2006; Han et al., 2008; Karanasios et al., 2010; Park et al., 2017; Kwiatek and Carman, 2020; Kwiatek et al., 2020). The key differences from fungi in the metabolic pathways that control membrane synthesis as well as emerging evidence for a tumor suppressor function of human CTDNEP1 motivate further analysis of the regulation of this pathway in mammalian cells. Loss of function mutations in CTDNEP1 are frequently observed in specific subtypes of medulloblastomas and were recently found to be enriched in medulloblastomas with MYC-amplification and genomic instability (Jones et al., 2012; Northcott et al., 2012; Luo et al., 2023). We had previously shown that ER expansion interferes with chromosome segregation causing higher rates of micronuclear formation (Merta et al., 2021), suggesting a link between dysregulated lipid synthesis and chromosomal instability in cancer cells. CTDNEP1 influences the lipid profile of the inner nuclear membrane (INM) as well as the stability of the inner nuclear envelope protein Sun2 through direct membrane sensing mechanism (Krshnan et al., 2022; Lee et al., 2023). CTDNEP1 has been implicated in regulating BMP signaling (Satow et al., 2006; Sakaguchi et al., 2013; Hayata et al., 2015; Darrigrand et al., 2020), which also involves nuclear envelope-associated proteins (Bengtsson, 2007). The relationship of CTDNEP1 to lipid synthesis, genomic instability, and the nuclear envelope underscores the importance of understanding how CTDNEP1 is regulated and how this regulation may be impacted in different cellular states.

We show that CTDNEP1 has evolved an N-terminal amphipathic helix required for its targeting to the ER/nuclear envelope and the surface of lipid droplets. The integral membrane binding partner, Nuclear Envelope Phosphatase 1 Regulatory subunit 1 (NEP1R1), temporarily protects CTDNEP1 from membrane extraction by the AAA+-ATPase p97 and degradation by the proteasome. Both peripheral membrane binding and stabilization of CTDNEP1 are required to promote lipin 1 dephosphorylation and restrict ER membrane production under normal conditions. In cells fed with excess fatty acids, CTDNEP1 phosphatase activity restricts lipid droplet formation; however, the same mechanism that stabilizes ER-associated CTDNEP1 to limit membrane production is no longer relevant to its role in lipid droplet biogenesis. Our work suggests that metabolic reprogramming from membrane synthesis to lipid storage rewires CTDNEP1 complex formation with NEP1R1 to locally control lipid droplet biogenesis. Our findings underscore the role of CTDNEP1 as a major regulator of lipid synthesis in the ER and highlight the importance of gaining a mechanistic understanding for how CTDNEP1 controls lipid homeostasis, especially in light of its critical role as a potential tumor suppressor.

## Results

### Different mechanisms target and stabilize CTDNEP1 at ER/nuclear membranes

CTDNEP1 is a member of the C-Terminal Domain-family of Ser/Thr phosphatases named after the founding member Scp1, which is the protein phosphatase for the CTD of RNA-PolII (Seifried et al., 2013). CTD-family phosphatases are part of an ancient superfamily of haloacid dehalogenases (HAD) defined by a signature DXDX(T/V) motif in the catalytic domain that carries out the phosphoryl transfer reaction (Figure 1A). CTD-family phosphatases evolved independently from PP1/PP2 alkaline phosphatases or tyrosine phosphatases and appear to have exquisite specificity for their substrates; however, little is known about how different members of the CTD-family of protein phosphatases recognize their distinct substrates (Seifried et al., 2013). Genetic studies in yeast uncovered Spo7 as a binding partner for Nem1 (yeast CTDNEP1; Siniossoglou et al., 1998) and later secondary structure analysis identified the integral membrane protein NEP1R1 (for Nuclear Envelope Phosphatase 1 Regulatory subunit 1, Spo7 in yeast; Han et al., 2012) as a Spo7 analogue and binding partner for human CTDNEP1 (Figure S1A-B, Figure 1A). CTDNEP1 is active towards a generic substrate *in vitro* (Kim et al., 2007), suggesting that NEP1R1 may not influence its mechanism of action, although whether this is the case in cells remains unknown. In human cells, transient expression of NEP1R1 and CTDNEP1 under control of constitutive promoters increased the protein levels of exogenous CTDNEP1 as well as lipin dephosphorylation (Han et al., 2012). The effect of NEP1R1 on the protein abundance, localization and activity of endogenous CTDNEP1 in cells had not been tested.

**Figure 1.**
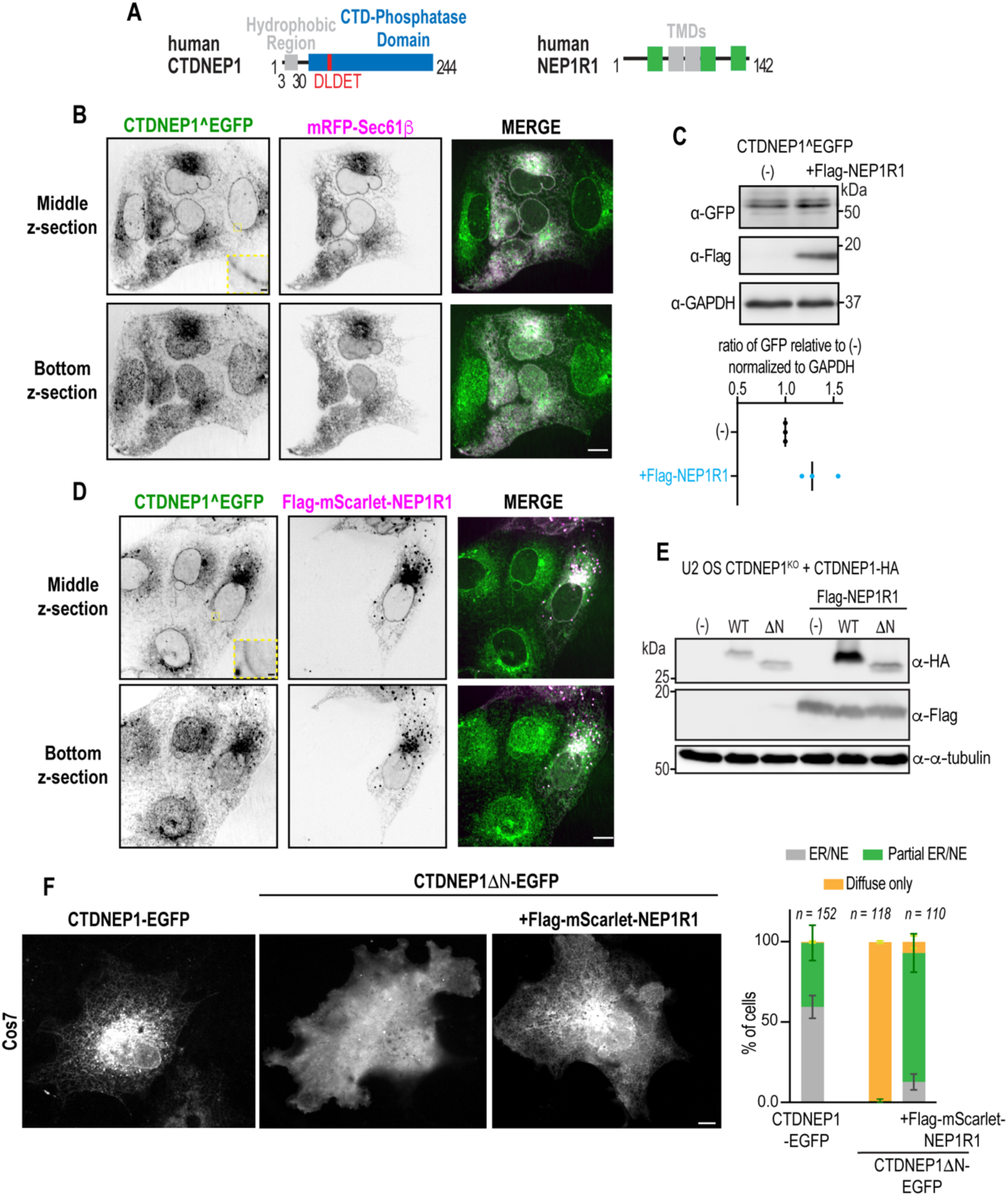
ER and nuclear envelope targeting and stabilization of CTDNEP1 by NEP1R1 depends on its N-terminal hydrophobic region. A) Schematic representation of CTDNEP1 and NEP1R1 protein domains. B) Representative spinning disk confocal images of CTDNEP1^EGFP and mRFP-Sec61β in living cells. Inset shows nuclear envelope localization of CTDNEP1^EGFP. Scale bar, 10µm; Scale bar inset, 1µm; N = 2 independent experiments. C) Top: Immunoblot of whole cell lysates from CTDNEP1^EGFP expressing cells transfected with Flag-NEP1R1. Bottom: Plot represents mean intensity value of CTDNEP1^EGFP relative to no transfection, normalized to GAPDH. Mean and N = 3 independent experiments shown. D) Representative spinning disk confocal images of CTDNEP1^EGFP in cells transfected with Flag-mScarlet-NEP1R1. Inset shows nuclear envelope localization of CTDNEP1^EGFP. Scale bar, 10µm; Scale bar inset, 1µm; N = 2 independent experiments. E) Immunoblot of whole cell lysates from CTDNEP1-deleted cells co-transfected with CTDNEP1-HA variants and Flag-NEP1R1. N = 3 independent experiments. F) Left: Representative spinning disk confocal images of indicated CTDNEP1-EGFP variants transiently transfected in living Cos-7 cells. Scale bar, 10µm. Right: Plot, blind quantification of CTDNEP1-EGFP localization under conditions on the left. Mean ± SD shown, n = individual cells counted. N = 3 independent experiments.

In our prior work, we generated a CRISPR-Cas9 gene edited cell line with EGFP inserted at the endogenous locus of CTDNEP1 (CTDNEP1^EGFP, Lee et al., 2023) to bypass difficulties in producing reliable antibodies that recognize endogenous CTDNEP1. Using this cell line, we show that CTDNEP1^EGFP localizes in a punctate pattern at the NE, similar to our prior reports (Lee et al., 2023), and that CTDNEP1 co-localizes with an ER marker in the perinuclear region (mRFP-Sec61β) and faintly to ER tubules (Figure 1B). We also observed CTDNEP1^EGFP puncta that did not colocalize with the ER marker and these often colocalized with the lysosomal marker Lamp1-mScarlet (Figure S1C). NEP1R1 overexpression increased endogenous CTDNEP1^EGFP protein levels by approximately 30% (Figure 1C). In the presence of NEP1R1, CTDNEP1 is more uniform throughout the ER and the NE (Figure 1D) and there is an increased abundance of puncta containing CTDNEP1 and NEP1R1 that resemble lysosomal structures (Figure 1D). RNAi-depletion of endogenous NEP1R1 decreased the ER/nuclear envelope subcellular localizations of CTDNEP1^EGFP, although punctate-like structures still remained, which may represent a more stable pool of CTDNEP1 (Figure S1D, S1E). The reduction in CTDNEP1^EGFP fluorescence signal in NEP1R1 RNAi-depleted cells was not because of a decrease in CTDNEP1 mRNA levels (Figure S1D) and was similar to, but to a lesser degree than RNAi-depletion of CTDNEP1 (Figure S1E). Together, these results indicated that NEP1R1 broadly effects the protein levels and ER/nuclear envelope localization of CTDNEP1.

Work in budding yeast showed that Nem1’s membrane association is mediated by both binding to Spo7 through its C-terminal catalytic domain and through its N-terminal hydrophobic helix (Siniossoglou et al., 1998). To test if the catalytic domain of CTDNEP1 is sufficient for targeting and stabilization of CTDNEP1 by NEP1R1, we transiently co-expressed NEP1R1 with full length CTDNEP1 or with its N-terminal hydrophobic region deleted (WT and ΔN, residues 3-30). NEP1R1 co-expression substantially increased the protein levels of WT CTDNEP1, but only slightly increased the protein levels of CTDNEP1ΔN (Figure 1E). Furthermore, overexpressed wild type CTDNEP1-EGFP uniformly localized to the ER/NE (Figure 1F), whereas CTDNEP1ΔN was diffusely localized in the cytoplasm (Figure 1F, Figure S1F). The ER/NE membrane localization of CTDNEP1 was only partially restored in cells transiently co-expressing CTDNEP1ΔN and NEP1R1 (Figure 1F). The fact that high expression of NEP1R1 was unable to stabilize or fully recruit CTDNEP1ΔN suggested that either the C-terminal phosphatase domain of CTDNEP1 is not sufficient to bind to NEP1R1 or membrane targeting of CTDNEP1 through its N-terminus is necessary for NEP1R1 binding.

### The N-terminus of CTDNEP1 encodes an amphipathic helix that is sufficient for targeting the ER, nuclear envelope and the surface of lipid droplets

We next characterized the N-terminal hydrophobic helix of CTDNEP1 to understand the relationship between membrane association and stabilization of CTDNEP1. The hydrophobic region in Nem1 is 39 amino acids, and prior reports have suggested that it is a double pass transmembrane domain (Siniossoglou et a*l.*, 1998; Figure S1A). Human CTDNEP1 contains a shorter hydrophobic helix of 28 amino acids (residues 3-30) that had been predicted to encode for a membrane spanning helix (Kim et al., 2007); however, with this orientation the C-terminal phosphatase domain would face the ER lumen (Figure S1G). Our *in-silico* Heliquest analysis and AlphaFold complex structure prediction suggested that the N-terminal residues 3-30 of CTDNEP1 encode for a membrane-associated amphipathic helix (Figure S1H, S1I). Amphipathic helices bind at the polar-non-polar interface of membranes (Drin and Antonny, 2010; Gimenez-Andres et al., 2018), which would fit with the expected cytoplasmic/nucleoplasmic facing orientation of the phosphatase domain of CTDNEP1 (Figure S1G, Lee et al., 2023). Multiple sequences alignments did not reveal high amino acid identity of the N-terminal region of CTDNEP1 across metazoans tested, however Heliquest predictions showed that the amphiphilicity of CTDNEP1’s N-terminus is conserved (Figure S2A, S2B).

Several lines of evidence further corroborated that the N-terminus of CTDNEP1 encodes for an amphipathic helix. The N-terminus (residues 3-30) fused to EGFP alone (hereafter AH-EGFP) localizes to ER/NE membranes when transiently overexpressed (Figure 2A). Mutating residues that preserve the positive charge of the polar face did not affect the NE/ER localization (AH(3R/K)-EGFP; Figure 2A). In contrast, mutating bulky hydrophobic residues (AH(2F/2L/A)-EGFP; Figure 2A) or replacing the positively charged residues (3RK) residues to aspartic acids (AH(3RK/D)-EGFP; Figure 2A) abolished ER/NE membrane targeting. Thus, the positively charged and bulky hydrophobic residues of the N-terminus of CTDNEP1 are necessary for its membrane targeting, which is in line with the *in silico* prediction that this region encodes for a peripheral membrane binding amphipathic helix.

**Figure 2.**
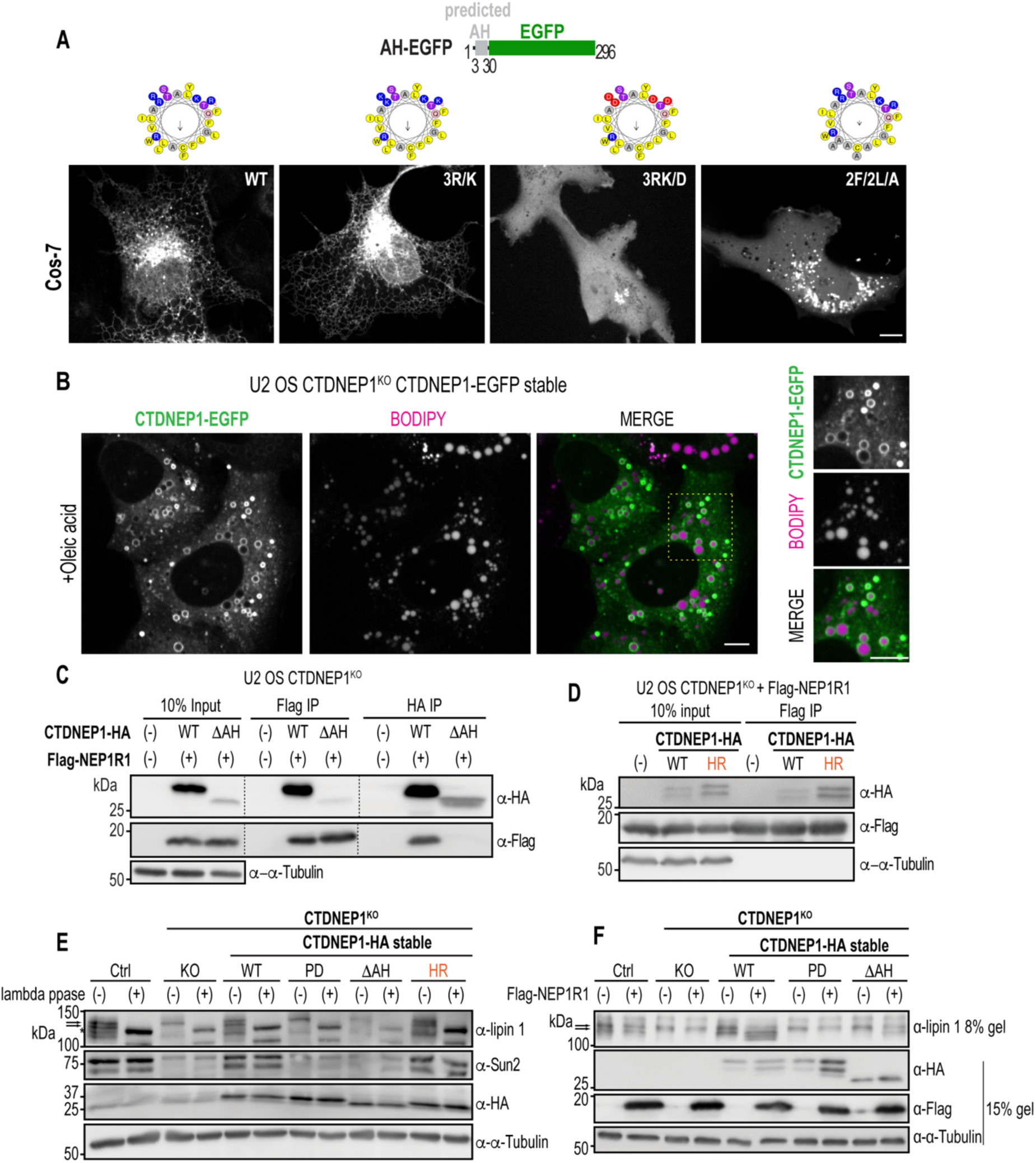
Membrane-associated CTDNEP1 binds NEP1R1 and is functional. A) Top: Schematic of the N-terminal amphipathic helix of CTDNEP1 fused to EGFP (AH-EGFP). Bottom: Representative spinning disk confocal images of AH-EGFP mutant variants transiently transfected in living Cos-7 cells. N = 2 (3R/K, 3RK/D and 2F/2L/A) or 3 (WT) independent experiments B) Representative spinning disk confocal images of stably expressed CTDNEP1-EGFP in living U2 OS CTDNEP1 deleted cells stained with Bodipy C12 and fed with 200µM Oleic acid for 24hrs. Insets show CTDNEP1-EGFP localization surrounding neutral lipid marker, Bodipy C12. N = 3 independent experiments C-D) Immunoblot of immunoprecipitated CTDNEP1-HA variants and Flag-NEP1R1 from cell lysates of U2 OS CTDNEP1-deleted cells transiently expressing indicated constructs. N = 2 independent experiments. E) Immunoblot of whole cell lysates treated with lambda phosphatase from CTDNEP1-deleted cells stably expressing CTDNEP1-HA variants. N = 2 independent experiments F) Immunoblot of whole cell lysates of CTDNEP1-deleted cells stably expressing CTDNEP1-HA variants transfected with Flag-NEP1R1. N = 2 independent experiments. For all: Scale bars, 10µm.

Selective permeabilization of the plasma membrane by digitonin showed that the C-terminus of CTDNEP1 faces the cytoplasm (Figure S2C). In addition, the predicted AH sequence tagged at both the N- and C-terminus localized to the ER/NE in digitonin-permeabilized cells (Figure S2D, S2E). These data provide further evidence that CTDNEP1 and its N-terminal predicted amphipathic helix do not cross the membrane.

Large bulky hydrophobic residues of amphipathic helices bind to packing defects present in the monolayer that surrounds fat filled lipid droplets (Prevost et al., 2018), so we tested if CTDNEP1 can localize to the surface of lipid droplets. In cells fed with exogenous fatty acids, CTDNEP1-EGFP localized to the rim enclosing a neutral lipid marker (Figure 2B) supporting the likelihood that CTDNEP1’s N-terminus is a membrane associated amphipathic helix. A CTDNEP1 chimera with its N-terminal amphipathic helix (aa 3-30) replaced with the *S. Cerevisiae* Nem1 hydrophobic region (aa 87-126) (Nem1HR-CTDNEP1) localized to ER/NE membranes (Figure S2F), but not to the surface of lipid droplets (Figure S2G), consistent with prior results that Nem1 localizes to the ER adjacent to, but not at the surface of, lipid droplets (Adeyo et al., 2011). We conclude that the N-terminus of CTDNEP1 has evolved to encode a peripheral membrane binding amphipathic helix critical for its targeting to ER, nuclear envelope and lipid droplets.

### ER/nuclear membrane targeting of CTDNEP1 promotes complex formation with NEP1R1 and lipin 1 dephosphorylation

We predicted that membrane targeting of CTDNEP1 is a prerequisite for complex formation with NEP1R1 and thus lipin 1 dephosphorylation. Reciprocal co-immunoprecipitation studies showed weak binding between CTDNEP1-HA without its N-terminal amphipathic helix (hereafter referred to as ΔAH) and Flag-NEP1R1 (Figure 2C), consistent with the subtle stabilization and only partial ER recruitment of ΔAH in cells overexpressing NEP1R1 (see Figure 1E, 1F). In contrast, the Nem1HR-CTDNEP1 chimeric protein that is constitutively localized to ER/nuclear membrane co-immunoprecipitated with Flag-NEP1R1 (Figure 2D), suggesting that membrane binding allows NEP1R1 complex formation. In cells stably expressing CTDNEP1 variants, wild type, but not phosphatase dead (PD) CTDNEP1, and the Nem1HR-CTDNEP1 chimera, promote lipin 1 dephosphorylation and Sun2 stability to a significantly greater extent than ΔAH (Figure 2E). Thus, CTDNEP1 must be membrane-bound to perform its catalytic function towards lipin 1. In line with this, ΔAH showed some activity towards lipin 1 dephosphorylation only in the presence of overexpressed Flag-NEP1R1 that partially recruits it to the ER/nuclear membranes (Figure 2F; see also Figure 1F). Overexpression of Flag-NEP1R1 greatly enhanced lipin 1 dephosphorylation by wild type CTDNEP1 (Figure 2F), as seen by the faster migrating bands that are also collapsed to a single band in cells treated with lambda phosphatase (Figure 2E), as previously shown (Han et al., 2012). Thus, both membrane targeting and CTDNEP1-NEP1R1 complex formation allow CTDNEP1 to perform its functions in dephosphorylation of lipin 1 and stabilization of Sun2.

### Membrane-bound CTDNEP1 is ubiquitylated and targeted for proteasomal degradation in a p97-dependent manner

The increase in CTDNEP1 protein levels in the presence of NEP1R1 prompted us to test if NEP1R1 protects CTDNEP1 from proteasomal degradation. Treating cells with cycloheximide to prevent protein translation revealed that wild type CTDNEP1-HA is short-lived (∼50% of the protein is degraded within 1 hr; Figure 3A, 3B) and that its degradation was slowed by a proteasome inhibitor MG132, indicating that it is a target of degradation by the proteasome (Figure 3A, 3C). In the absence of its N-terminal ER/NE targeting region, CTDNEP1ΔAH was rapidly degraded by the proteasome (∼81% of existing protein remained in 1 hour; Figure 3A-3C). These data indicate that both soluble and membrane-bound CTDNEP1 are targeted for proteasomal degradation.

**Figure 3.**
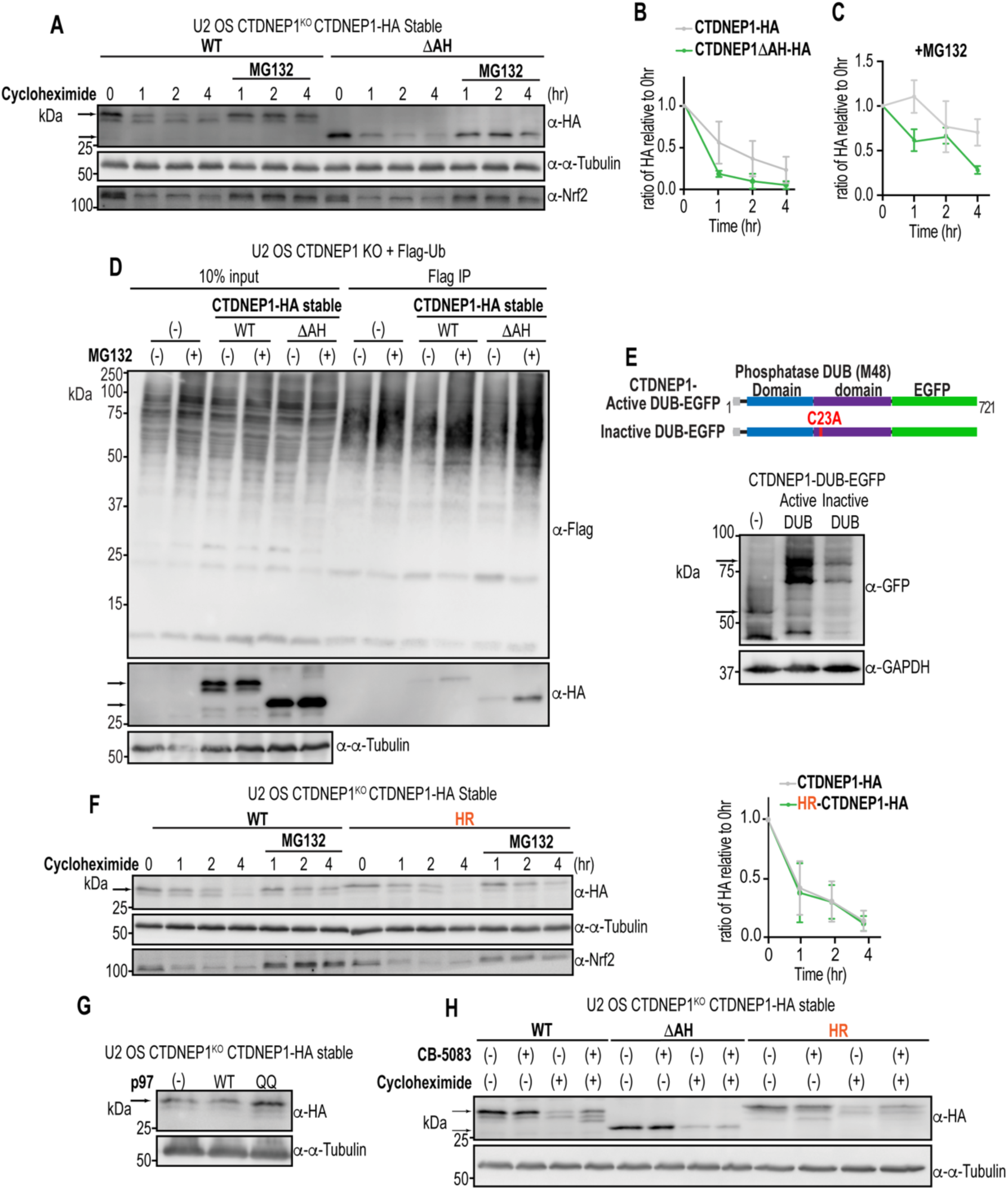
Membrane-bound CTDNEP1 is targets for degradation by the ubiquitin-proteasome system in a p97-dependent manner. A) Immunoblot of whole cell lysates of CTDNEP1 deleted cells stably expressing CTDNEP1-HA variants treated with cycloheximide ± MG132 for the indicated times. B-C) Plot represents mean intensity value of top band of CTDNEP1-HA relative to its initial timepoint (0hr) normalized to tubulin in either (B) Cycloheximide or (C) Cycloheximide + MG132 conditions. Mean ± SD shown. N = 3 (Cycloheximide + MG132) or 4 (Cycloheximide) independent experiments. D) Immunoblot of immunoprecipitated CTDNEP1-HA with Flag-Ubiquitin in U2 OS CTDNEP1-deleted cells stably expressing CTDNEP1 variants, transiently expressing Flag-Ubiquitin and treated with MG132 for 2 hrs. N = 2 independent experiments. E) Top: Schematic of CTDNEP1 fused to M48 deubiquitylase domain (DUB) and EGFP. Bottom: Immunoblot of whole cell lysates of U2 OS cells transfected with the indicated constructs. N = 2 independent experiments. F) Left: Immunoblot of whole cell lysates of CTDNEP1-deleted cells stably expressing CTDNEP1-HA variants and treated with cycloheximide ± MG132 for the indicated time course; Right: Plot represents mean intensity value of top band of CTDNEP1-HA relative to its initial timepoint (0 hr) normalized to tubulin. Mean ± SD shown. N = 3 independent experiments. G) Immunoblot of whole cell lysates of CTDNEP1-deleted cells stably expressing CTDNEP1-HA transfected with p97 variants. N = 2 independent experiments. H) Immunoblot of whole cell lysates of CTDNEP1-deleted cells stably expressing CTDNEP1-HA variants and treated with cycloheximide and/or CB-5083 to inhibit p97 for 4hrs. N = 3 independent experiments.

Prior large-scale screens identified direct ubiquitylation sites in CTDNEP1 (Rose et al., 2016). In cells treated with a proteasomal inhibitor to increase ubiquitylated proteins, full length and ΔAH CTDNEP1 co-immunoprecipitate with Flag-Ubiquitin (Figure 3D), providing supportive evidence that ubiquitylation of CTDNEP1 targets it for proteasomal degradation. The relatively increased amount of ΔAH that pulled down with Flag-Ubiquitin suggested that non-membrane bound CTDNEP1 may be more prone to ubiquitylation and degradation (Figure 3D). Fusion of full length CTDNEP1 to a deubiquitylating domain that cleaves polyubiquitylated chains increased the protein levels of CTDNEP1 and this was dependent on the deubiquitylase catalytic activity (Figure 3E; Schlieker et al., 2007). The fusion protein accumulated in both the cytoplasm and ER/NE (Figure S3A), suggesting that CTDNEP1 protein stability depends on polyubiquitylation. The constitutively membrane bound Nem1HR CTDNEP1 chimera (Figure 3F) and CTDNEP1-PD (Figure S3B) are degraded at a similar rate as wild type CTDNEP1. Thus, membrane-associated CTDNEP1 is a target of proteasomal degradation and this is independent of its catalytic active site.

To visually observe the short-lived subcellular pools of CTDNEP1 protein, we imaged CTDNEP1 knockout cells stably expressing wild type CTDNEP1-EGFP, which is expressed at higher levels than endogenously tagged CTDNEP1^EGFP (Figure S3C, S3D). Cycloheximide treatment showed uniform loss of EGFP signal from the ER and NE (Figure S3C). In cells additionally treated with MG132, CTDNEP1-EGFP accumulated to punctate structures in the perinuclear region, the majority of which co-localized with the lysosomal marker Lamp1-mScarlet (Figure S3C, S3E). These data suggest that inhibition of the proteasome results in existing pools of CTDNEP1 to traffic to the lysosome, which may prevent its aberrant accumulation at the ER/NE. We tested if CTDNEP1 is degraded by the lysosome by treating cells expressing CTDNEP1-HA with lysosomal inhibitor Bafilomycin A1 (BafA1, Yoshimori et al., 1991) (Figure S3F). BafA1 treatment results in a mild increase in levels of total CTDNEP1 protein suggesting that a small pool of CTDNEP1 is degraded by the lysosome, in line with its co-localization with the lysosomal marker (see Figure S1C, S3C). BafA1 treatment did not prevent the rapid loss of existing CTDNEP1-HA protein in cycloheximide-treated cells (Figure S3F), suggesting that CTDNEP1 stability is mainly regulated by proteasome-mediated degradation, and trafficking to the lysosome may contribute to the degradation of excess membrane-bound CTDNEP1.

We next tested if the AAA^+^ ATPase VCP/p97 (Cdc48 in *S. cerevisiae*) protein complex that extracts proteins from the ER/NE and cooperates with the proteasome in the ER-associated protein degradation (ERAD) pathway participates in degradation of CTDNEP1. Overexpression of a dominant negative variant of p97 (p97 QQ) (Tsai et al., 2016) that no longer hydrolyzes ATP increased the protein levels of CTDNEP1-HA stably expressed in CTDNEP1 knockout cells (Figure 3G). p97 inhibition, either by overexpression of the p97 QQ dominant negative construct or by treating cells with a small molecule ATP competitive inhibitor of a domain of p97, also stabilized endogenous CTDNEP1-mAID-HA in DLD-1 cells (Figure S3G, Lee et al., 2023). Thus, p97 activity is required to promote degradation of CTDNEP1. Small molecule inhibition of p97 stabilized existing protein pools of wild type CTDNEP1 and the Nem1HR-CTDNEP1 chimera, but not soluble ΔAH CTDNEP1 (Figure 3H). Thus, a p97-dependent activity is required for the regulated degradation of membrane-bound CTDNEP1. Interestingly, NEP1R1 is long-lived and not degraded by the proteasome in the same time scale as CTDNEP1 (Figure S3H), suggesting that CTDNEP1, and not the CTDNEP1-NEP1R1 complex, is selectively targeted for proteasomal degradation.

Together, these data show that CTDNEP1 is regulated by multiple degradation pathways. The majority of endogenous CTDNEP1 is bound to ER/nuclear membranes, which promotes its function in lipin 1 dephosphorylation. This pool of CTDNEP1 is short-lived and degraded in a p97 and proteasomal-dependent manner.

### CTDNEP1 and NEP1R1 associate directly through a hydrophobic binding interface

Our data showing that CTDNEP1 protein levels correlate with those of NEP1R1 suggested that CTDNEP1-NEP1R1 complex formation may control the rate of proteasomal degradation of CTDNEP1. To test this possibility, we sought out to create mutant versions of either NEP1R1 or CTDNEP1 that disrupt complex formation without disrupting their overall domain architecture and targeting to the ER/NE. AlphaFold 3D modeling of the CTDNEP1/NEP1R1 heterodimer revealed a hydrophobic interface between the proteins consisting of mainly three residues in CTDNEP1 (CTDNEP1 M220, A223, V233) and two residues in NEP1R1 (NEP1R1 F30, L120) with hydrophobic amino acid side chains at its core (Figure 4A, 4B). Mutating the bulky, centrally positioned phenylalanine residue (F30) of NEP1R1 to either a small hydrophobic amino acid (F30A) or a charged amino acid (F30E) strongly reduced binding with CTDNEP1-HA, as shown by reciprocal co-immunoprecipitation experiments (Figure 4C). Variants of CTDNEP1-HA in which the hydrophobic V233 was mutated to a charged residue (V233E) or both M220/V233 were mutated to alanine also strongly reduced complex formation with Flag-NEP1R1 (Figure 4D). The ER/NE localization of the mutant versions of either NEP1R1 (Figure S4A) or CTDNEP1 (Figure S4B) was unaffected compared to the wild type tagged proteins, indicating that a change in subcellular localization does not underlie the lack of complex formation under these conditions.

**Figure 4.**
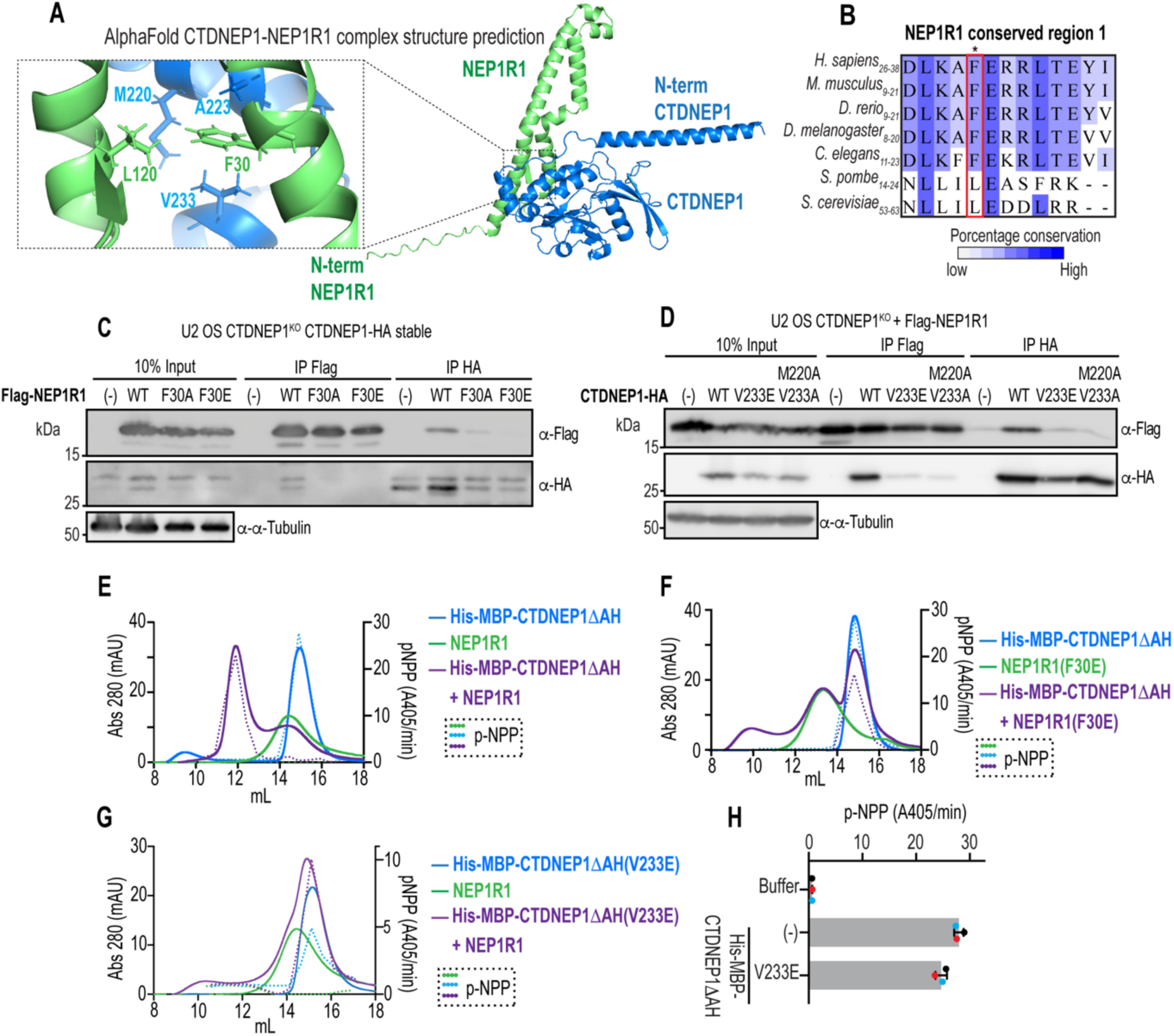
CTDNEP1 to NEP1R1 interaction *in vitro* and *in vivo* depends on a hydrophobic interface. A) AlphaFold structure prediction of the CTDNEP1-NEP1R1 complex. Zoom-in highlights residues at a binding interface between the proteins. B) Multiple sequence alignment of NEP1R1 amino acid residues at binding interface with CTDNEP1 based on (A). The metazoan-specific conserved phenylalanine residue is highlighted in a red box. C) Immunoblot of immunoprecipitated Flag-NEP1R1 variants with CTDNEP1-HA in U2 OS CTDNEP1-deleted cells stably expressing CTDNEP1-HA and transfected with mutant variants. N = 2 independent experiments. D) Immunoblot of immunoprecipitated CTDNEP1-HA mutant variants with Flag-NEP1R1 in U2 OS CTDNEP1 deleted cells transiently expressing indicated proteins. N = 2 independent experiments. E-G) Size exclusion chromatography of purified His6-MBP-CTDNEP1ΔAH and NEP1R1 variants. Solid lines are the elution profiles of the proteins and dotted lines are the pNPP assay results. H) p-NPP assay of His6-MBP-CTDNEP1ΔAH variants. Mean ± SD shown. N = 3 independent experiments.

We next determined if the hydrophobic interfaces disrupt direct binding between CTDNEP1 and NEP1R1 *in vitro* (see Figure 4A). Size exclusion chromatography of *in vitro* purified CTDNEP1ΔAH (tagged with His-MBP) combined with NEP1R1 co-eluted in peak fractions that were shifted relative to CTDNEP1ΔAH or NEP1R1 alone (Figure 4E), indicating that these proteins form a complex. CTDNEP1ΔAH phosphatase activity towards the generic substrate p-NPP confirmed co-elution of CTDNEP1 with wild type NEP1R1 (dotted lines in Figure 4E). The peak representing complex formation was absent when the NEP1R1(F30E) mutant was combined with CTDNEP1ΔAH (Figure 4F). Combining active CTDNEP1ΔAH(V233E) (tagged with His-MBP) with wild type NEP1R1 also did not result in the shifted peak that represents complex formation (Figure 4G), further verifying that the predicted hydrophobic interface formed by these residues facilitates complex formation. Quantitative comparison of the phosphatase activity of *in vitro* purified the CTDNEP1ΔAH(V233E) mutant towards a generic substrate demonstrated that this mutation does not disrupt the catalytic site of CTDNEP1 *in vitro* (Figure 4H). Thus, our *in vitro* and *in vivo* data identify key residues that facilitate CTDNEP1/NEP1R1 complex formation.

### NEP1R1 limits the rate of proteasomal degradation of CTDNEP1

The CTDNEP1 and NEP1R1 mutant variants that localize to the ER/nuclear envelope but are unable to heterodimerize allowed us to directly test how complex formation influences the stability and functions of CTDNEP1. We first overexpressed wild type Flag-NEP1R1 and found that the majority of existing CTDNEP1-HA protein was stable after 1 hour (Figure 5A, 5B; ∼70% of the protein remains within 1 hr). In cells overexpressing the NEP1R1(F30E) less than half of existing CTDNEP1-HA remained (Figure 5A, 5B), which is similar to the rate of degradation of CTDNEP1-HA in cells that were not overexpressing NEP1R1 (see Figure 3A). Overexpression of the NEP1R1(F30E) mutant did not increase endogenous CTDNEP1^EGFP protein levels (Figure S4C compared to Figure 1C). Interestingly, wild type Flag-NEP1R1 was long-lived regardless of heterodimer formation (Figure 5A). Moreover, while complex formation with NEP1R1 slows the rate of CTDNEP1 degradation, CTDNEP1 remains a highly regulated target of the proteasome in the presence of overexpressed wild type NEP1R1, as shown by the eventual loss of existing CTDNEP1-HA protein (Figure 5A) and its co-immunoprecipitation with Flag-ubiquitin upon inhibition of the proteasome (Figure 5C). The relative greater increase in band intensity of CTDNEP1 co-immunoprecipitated with Flag-ubiquitin in cells treated with a proteasomal inhibitor compared to NEP1R1 suggests that ubiquitylated CTDNEP1 accumulates to a greater extent than NEP1R1 (Figure 5C). Thus, CTDNEP1 is highly prone to ubiquitylation and degradation while NEP1R1 is relatively more stable. We conclude that direct binding of NEP1R1 to CTDNEP1 stabilizes CTDNEP1 by limiting the rate of its proteasomal degradation.

**Figure 5.**
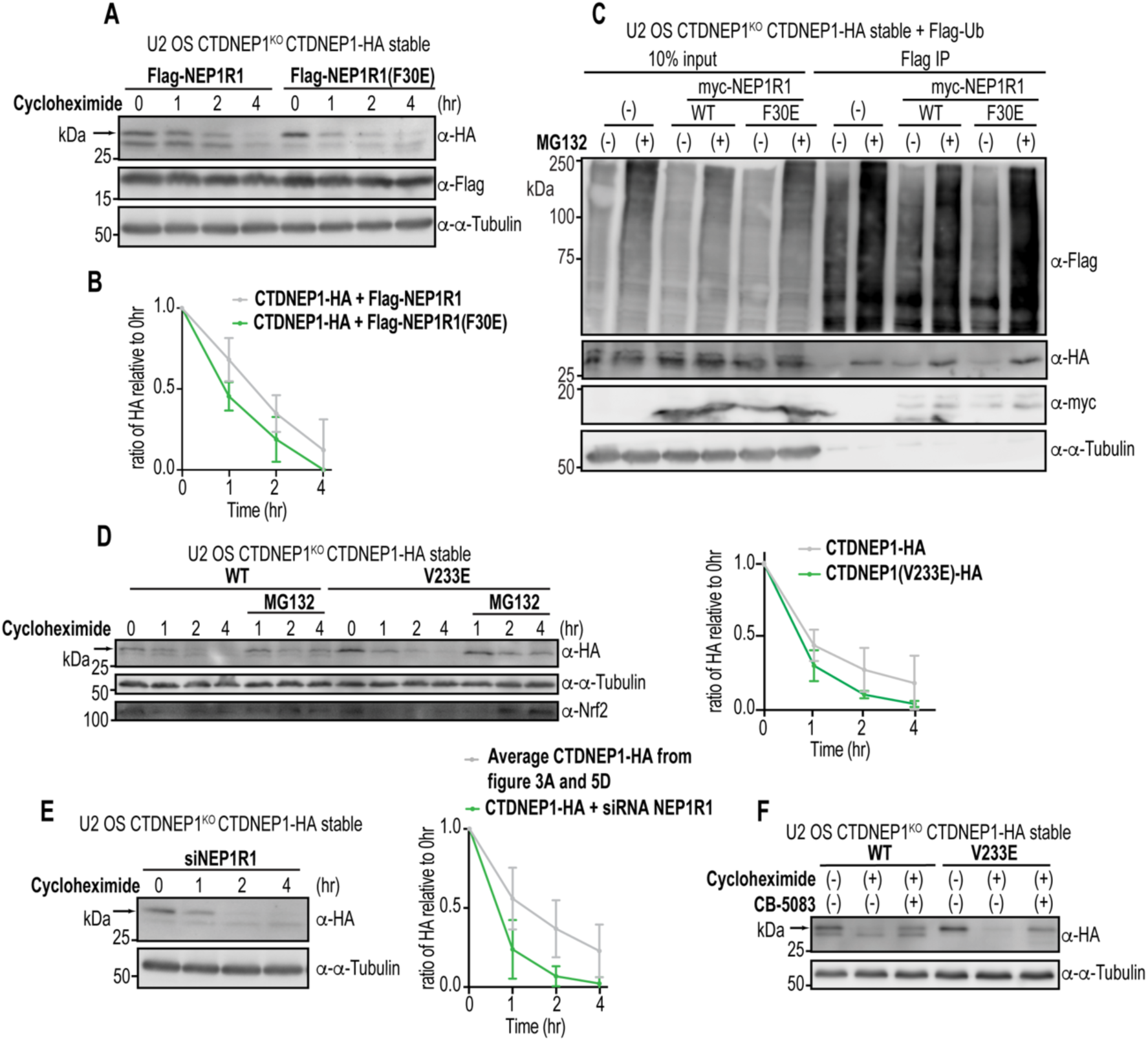
NEP1R1 is relatively long-lived and limits the rate of CTDNEP1 protein degradation. A) Immunoblot of whole cell lysates of CTDNEP1-deleted cells stably expressing CTDNEP1-HA transfected with Flag-NEP1R1 variants and treated with cycloheximide for the indicated time course. B) Plot represents mean intensity value of top band of CTDNEP1-HA relative to its initial timepoint (0 hr) normalized to tubulin. Mean ± SD shown. N = 3 independent experiments. C) Immunoblot of immunoprecipitated Flag-Ubiquitin with myc-NEP1R1 wild type and F30 mutant variant from U2 OS CTDNEP1-deleted cell lysates stably expressing CTDNEP1-HA and treated with MG132 for 4hrs. N = 2 independent experiments. D) Left: Immunoblot of whole cell lysates of CTDNEP1-deleted cells stably expressing CTDNEP1-HA mutant variants and treated with cycloheximide ± MG132 for the indicated time points. Right: Plot represents mean intensity value of top band of CTDNEP1-HA relative to its initial timepoint (0 hr) normalized to tubulin. Mean ± SD shown. N = 3 independent experiments. E) Left: Immunoblot of whole cell lysates of CTDNEP1-deleted cells stably expressing CTDNEP1-HA, RNAi-depleted of endogenous NEP1R1 and treated with cycloheximide for indicated time points. Right: Plot represents mean intensity value of top band of CTDNEP1-HA relative to its initial timepoint (0 hr) normalized to tubulin. Mean ± SD shown. N = 3 (RNAi NEP1R1) or 7 (CTDNEP1-HA from 3A and 5D) independent experiments. F) Immunoblot of whole cell lysates of CTDNEP1-deleted cells stably expressing CTDNEP1-HA variants treated with cycloheximide and/or CB-5083 to inhibit p97 for 4hrs. N = 3 independent experiments.

We next assessed the degradation rate of CTDNEP1 in cells RNAi-depleted for NEP1R1 and of the CTDNEP1(V233E) mutant that is significantly reduced in its binding to NEP1R1 (see Figure 4D). In both cases, CTDNEP1 not bound to NEP1R1 is very short-lived (only ∼30% of the protein remains within 1 hr; Figure 5D, 5E), further confirming the effect of NEP1R1 on limiting the degradation rate of CTDNEP1. Furthermore, p97 activity is required for degradation of existing CTDNEP1(V233E)-HA protein (Figure 5F), suggesting that p97 extracts CTDNEP1 from membranes even when it is not bound to NEP1R1. Together, our data suggest that NEP1R1 acts as an ER/NE membrane scaffold that temporarily shelters membrane-bound CTDNEP1 from proteasomal degradation.

### NEP1R1 regulates the stability of CTDNEP1 to restrict ER membrane expansion

CTDNEP1 activity is required to limit ER membrane biogenesis through lipin 1 dephosphorylation (Merta et al., 2021). While our data show that membrane association of CTDNEP1 and NEP1R1 overexpression, which increases CTDNEP1 protein, promote lipin 1 dephosphorylation, it remains unclear if this function relies on direct binding of CTDNEP1 with NEP1R1. The CTDNEP1(V233E)-HA mutant that is active *in vitro* and localizes to ER-nuclear membranes but does not bind to NEP1R1 (see Figure 4D, 4H and S4B) provides a tool to test if CTDNEP1-NEP1R1 binding is required for lipin 1 dephosphorylation. In CTDNEP1 knockout cells stably expressing CTDNEP1(V233E), lipin 1 is not dephosphorylated to the same extent as cells expressing wild type CTDNEP1 or the Nem1HR CTDNEP1 chimera (Figure 6A). In line with the reduction in lipin 1 dephosphorylation, Sun2 levels are not stabilized with the CTDNEP1(V233E) mutant (Figure 6A). Furthermore, the CTDNEP1ΔAH and V233E mutants do not support nuclear translocation of lipin1β-GFP upon inhibition of mTORC1 (Figure S5A, S5B). These results demonstrate that CTDNEP1-NEP1R1 complex formation is necessary for dephosphorylation of lipin 1, and thus nuclear translocation of lipin 1 and Sun2 stability. Expression from a constitutive promoter increases the steady state protein levels of the V233E mutant, but this is not sufficient for lipin 1 dephosphorylation and Sun2 stability. Thus, in addition to protection from proteasomal degradation, NEP1R1 binding likely promotes CTDNEP1 activities through other unknown mechanisms.

**Figure 6.**
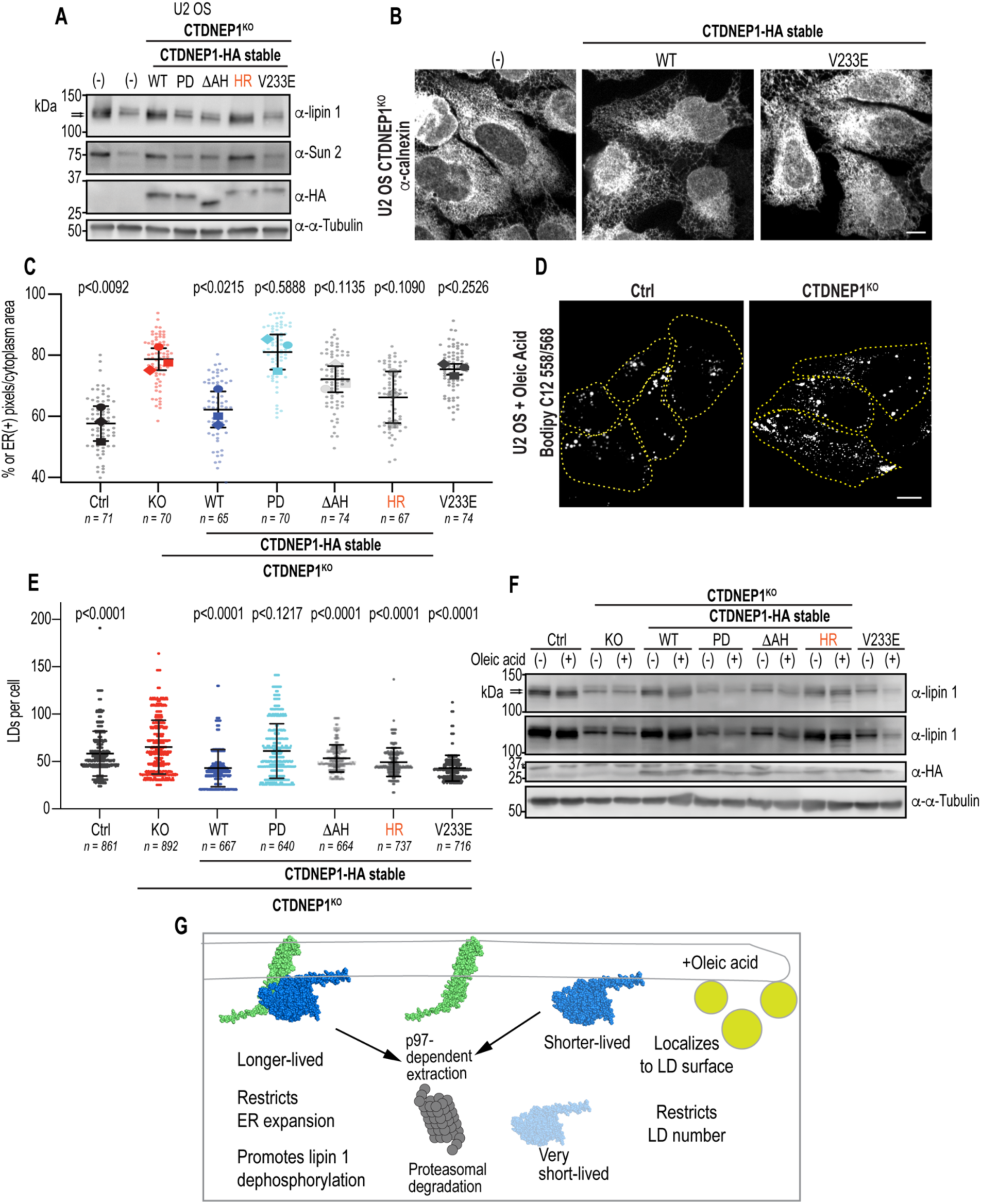
CTDNEP1 complex formation with NEP1R1 restricts ER membrane expansion but is not required for CTDNEP1’s function in limiting lipid droplet biogenesis. A) Immunoblot of whole cell lysates of CTDNEP1 deleted cells stably expressing CTDNEP1-HA mutant variants. N = 3 independent experiments. B) Representative spinning disk confocal images of calnexin immunostaining in fixed U2 OS CTDNEP1-deleted cells stably expressing CTDNEP1-HA variants. C) Plot represents percent area of calnexin fluorescence signal (cell area subtracted by nuclear area) for cells from indicated cells. Individual data points and mean ± SD shown. N = 3 independent experiments; p-values are paired t tests of replicate mean; all p values are relative to U2 OS CTDNEP1-deleted cells. D) Representative spinning disk confocal images of Bodipy C12 staining in living U2 OS and U2 OS CTDNEP1-deleted cells treated with oleic acid for 24 hours. Dotted lines indicate the edges of the cells. E) Plot represents quantification of the number of lipid droplets per cell in the indicated cells using semi-automated algorithm. Mean ± SD shown. N = 4 (Ctrl and CTDNEP1 deleted cells) or 5 (CTDNEP1-HA stable variants) independent experiments. p values, One-way Anova with Dunnet’s multiple comparisons test, all p values are relative to CTDNEP1-deleted cells. F) Immunoblot of whole cell lysates of CTDNEP1-deleted cells stably expressing CTDNEP1-HA mutant variants treated with Oleic acid for 24hrs. N = 2 independent experiments. G) Schematic representing different pools of CTDNEP1 and their relative stabilities and roles in lipid synthesis. For all: Scale bars, 10µm. n = individual cells counted.

The extent at which different mutant variants of CTDNEP1 promote lipin 1 dephosphorylation correlated with the ability to rescue the increase in ER size and defects in nuclear solidity resulting from loss of CTDNEP1 (Figure 6B, 6C, S5C-S5E). Thus, complex formation is necessary for CTDNEP1 to restrict ER membrane biogenesis through lipin 1 regulation. In line with this, siRNA depletion of endogenous NEP1R1 results in ER expansion (Figure S5F-S5H), and this is rescued by expression of siRNA resistant wild type, but not F30E Flag-NEP1R1 (Figure S5F-S5H). We conclude that binding to NEP1R1 is essential for CTDNEP1 to perform its function in regulating ER membrane biogenesis in cells.

### CTDNEP1 controls lipid droplet number independent of NEP1R1 binding

The mutant version of CTDNEP1 that no longer binds NEP1R1 provided us with a tool to determine the effect of disrupting complex formation in cells fed excess fatty acids. Oleic acid promotes dephosphorylation of lipin 1 and translocation to membranes (Harris et al., 2007; Kim et al., 2007). It was previously shown that in some cell types overexpression of CTDNEP1 alone did not completely promote dephosphorylation of overexpressed lipin 1 (Kim et al., 2007). In budding yeast, Nem1 (human CTDNEP1), Spo7 (human NEP1R1) and Pah1 (human lipin) are required for TAG synthesis - deletion of any of these genes leads to ER membrane expansion and reduced lipid droplet number (Siniossoglou et al., 1998; Santos-Rosa et al., 2005; Kim et al., 2007; Adeyo et al., 2011; Han et al., 2012; Papagiannidis et al., 2021;). Furthermore, overexpression of human CTDNEP1 with NEP1R1, but not overexpression of CTDNEP1 alone, restored reduced LD and TAG levels resulting from *spo7Δ* or *nem1Δspo7Δ* mutants (Han et al., 2012). These data had suggested that complex formation between these proteins is required for lipid droplet biogenesis, especially in budding yeast where most of these studies were conducted.

To test if CTDNEP1 has a role in LD formation in mammalian cells, we treated CTDNEP1 knockout cells with oleic acid to induce lipid droplet formation. CTDNEP1 knockout cells contained a greater number of lipid droplets per cell, and this was dependent on its phosphatase active site (Figure 6D, 6E). Surprisingly, expression of WT CTDNEP1 and all of the mutant variants (Nem1HR, ΔAH and V233E), except for phosphatase dead (PD), CTDNEP1 restored the increase in lipid droplet number resulting from loss of CTDNEP1 (Figure 6E). Interestingly, the CTDNEP1(V233E) mutant that is significantly reduced in NEP1R1 binding rescued lipid droplet number to the same extent as wild type CTDNEP1 levels (p < 0.9999 relative to WT CTDNEP1; see Figure 6E), whereas the Nem1HR and ΔAH variants restored lipid droplet number to a lesser extent (p < 0.0001 relative to WT CTDNEP1 for both; see Figure 6E). Furthermore, we observed the appearance of a faint faster migrating endogenous lipin 1 species induced by oleic acid treatment in all conditions except for CTDNEP1 knockout cells and CTDNEP1-PD expressing cells (Figure 6F). This suggested that NEP1R1 binding and membrane association are not required for oleic-acid stimulated dephosphorylation of lipin 1. The protein levels of lipin 1 are lower in the ΔAH and V233E mutant conditions (Figure 6F) suggesting that the effect of loss of CTDNEP1 on increased lipin 1 turnover may be through a distinct mechanism that still requires membrane and NEP1R1 binding. Together, these results indicate that CTDNEP1 limits the number of lipid droplets per cell under conditions of excess oleic acid, however this function is independent of NEP1R1 binding and partially dependent on its amphipathic helix.

In sum, ER/nuclear membrane targeting and NEP1R1 binding of CTDNEP1 are required for limiting ER membrane synthesis and promoting lipin 1 dephosphorylation and Sun2 stability. CTDNEP1 also restricts lipid droplet formation; however, this function does not require NEP1R1 binding. We conclude that under conditions of excess fatty acids CTDNEP1 function is rewired to restrict lipid droplet biogenesis providing evidence for context-dependent regulation and function of CTDNEP1.

## Discussion

We demonstrate that different mechanisms are involved in the targeting and stabilization of CTDNEP1 at ER/nuclear membranes. These mechanisms coordinately regulate CTDNEP1 to promote lipin 1 dephosphorylation for limiting ER membrane expansion. An N-terminal amphipathic helix in CTDNEP1 targets it to the ER and nuclear envelope where it is either rapidly degraded by the proteasome or is temporarily shielded from proteasomal degradation by NEP1R1- binding (Figure 6G). We identify a key interaction interface in CTDNEP1 and NEP1R1 that facilitates complex formation and show the importance of NEP1R1 binding in promoting the phosphatase activity of CTDNEP1 towards lipin 1 to counteract mTORC1-mediated cytoplasmic retention of lipin 1 in cells. Interestingly, CTDNEP1 activity also plays a role in limiting oleic-acid induced lipid droplet biogenesis, but its mode of regulation is no longer dependent on NEP1R1 binding. Thus, by dissecting how CTDNEP1 is regulated to restrict ER membrane biogenesis, we discovered that its regulation is rewired in the presence of excess fatty acids, which may control a pool of lipin 1 or other substrates important for lipid droplet biogenesis.

Work in yeast as elucidated the dependence of Nem1 on Spo7 in regulation of Pah1 and the effect of Pah1 dephosphorylation on its activity and membrane association (Siniossoglou et al., 1998; Karanasios et al., 2010; Han et al., 2012; Karanasios et al., 2013; Mirheydari et al., 2020; Jog et al., 2023). However, there are key differences for mammalian cells. In yeast, dephosphorylation of Pah1 by the Nem1/Spo7 complex promotes Pah1’s membrane association where it can readily dephosphorylate phosphatidic acid to DAG for TAG production and lipid droplet biogenesis (Santos-Rosa et al., 2005; O’Hara et al., 2006; Adeyo et al., 2011; Karanasios et al., 2013). In the absence of *nem1/spo7* or in yeast deleted of *pah1*, there is an increase in PA levels that feeds into membrane synthesis (Santos-Rosa et al., 2005; Papagiannidis et al., 2021). In mammalian cells, DAG is the precursor to both membrane and TAG synthesis (Stevenson et al., 2016). Increased PA in the presence of excess fatty acids can promote lipin 1’s membrane association and overcome its decreased levels or increased phosphorylation under most conditions (Eaton et al., 2013). In the absence of CTDNEP1, a dephosphorylated lipin 1 product is absent and there are lower levels of lipin 1; however fatty acid synthesis is upregulated, which may allow for lipin 1 to associate with membranes to maintain DAG production for membrane biogenesis (Merta et al., 2021).

We show that NEP1R1 binding is not required for CTDNEP1 to limit LD number in the presence of oleic acid. We also show that CTDNEP1 associates with NEP1R1 to promote lipin 1 dephosphorylation under normal conditions, and the loss of this regulation increases membrane biogenesis. Feeding cells excess fatty acids rewires CTDNEP1 to no longer require NEP1R1 binding for restricting LD formation. Interestingly, in yeast Nem1 localizes to ER-LD contact sites where it promotes local DAG production by Pah1 (Adeyo et al., 2011). Local DAG production plays a role in lipid droplet formation independent of TAG production (Choudhary et al., 2020). We show that CTDNEP1 localizes to the surface of lipid droplets. Replacing the amphipathic helix in CTDNEP1 with the Nem1 hydrophobic region prevents LD association. Thus, CTDNEP1 may have evolved an amphipathic helix that under conditions of LD formation allow its segregation from NEP1R1 where it can perform other local functions. It is interesting that in the presence of oleic acid only the V233E mutant fully restores the increase in LD number resulting from loss of CTDNEP1, whereas deletion of the amphipathic helix of CTDNEP1 and the Nem1HR chimera only partially restore LD formation. This suggests that a local role for CTDNEP1 at the surface of LDs, which may involve lipin 1 and possibly other substrates, is NEP1R1-independent.

CTDNEP1/NEP1R1 complex formation may be negatively regulated to promote a pool of CTDNEP1 to localize to LDs in the presence of oleic acid. In yeast, several post-translational events as well as protein binding partners have been shown to disrupt formation of the Nem1/Spo7 complex (Su et al., 2018; Dey et al., 2019; Papagiannidis et al., 2021; Saik et al., 2023;). Negative regulation of CTDNEP1/NEP1R1 complex formation and/or function may be related to the switch from membrane biogenesis to lipid storage in the presence of exogenous fatty acids.

Increasing evidence shows that crosstalk between the ER-associated ubiquitin proteasome pathway and regulation of lipid synthesis maintains ER lipid homeostasis (Stevenson et al., 2016; Olzmann and Carvalho, 2019). We show that membrane-bound CTDNEP1 is degraded through mechanisms that resemble ERAD pathway (Christianson et al., 2023). NEP1R1 is relatively stable at the ER and acts as a membrane scaffold to reduce the rate of proteasomal degradation of CTDNEP1. The fact that CTDNEP1 degradation is still active even with high levels of NEP1R1 suggests that the association between these proteins is regulated to release CTDNEP1 for targeting to the proteasome. CTDNEP1 is also degraded by lysosomes suggesting potential harmful effects of high levels of CTDNEP1 in cells. Future work is needed to determine the mechanism of CTDNEP1 ubiquitylation and targeting to the proteasome and how NEP1R1 binding limits proteasomal degradation of CTDNEP1.

CTDNEP1 and its metazoan homologues are nuclear-envelope localized and participate in many nuclear envelope-dependent processes through lipin regulation (Han et al., 2012; Bahmanyar et al., 2014; Penfield et al., 2020; Merta et al 2021., Lee et al., 2023). The combined effect of cytoplasmic enrichment of kinases that target lipin and nuclear envelope enrichment of CTDNEP1 may make nuclear-enriched lipin more active for DAG production at the inner nuclear membrane than in the cytoplasm/ER (Peterson et al., 2011; Merta et al., 2021; Lee et al., 2023). Whether there are differences in regulation of the complex at the nuclear envelope relative to the rest of the ER would provide insight into how DAG production at the inner nuclear membrane is regulated and may inform on the different roles of DAG in nuclear envelope functions.

The role of CTDNEP1 in counteracting mTOR-mediated lipogenesis and the emergence of CTDNEP1 as a tumor suppressor underscores the importance of understanding how CTDNEP1 performs its functions in regulating lipid synthesis. CTDNEP1 is also implicated in BMP signaling and in regulation of neural tube development suggesting potential context-dependent differences in its function (Satow et al., 2006). Moreover, metabolic disorders associated with lipin deficiency motivate further understanding of its regulation under different environmental conditions (Reue, 2009). Finally, the role of CTDNEP1 in regulation of lipid synthesis is critical to gain a broad understanding to how cells control their membrane content to accommodate different cellular functions.

## Acknowledgments

We thank M. Hochstrasser, J.M. Gendron, C. Schlieker (Yale University) for helpful discussions; Members of C. Schlieker, J.M. Gendron J. Bewersdorf, D. K. Breslow and P. De Camili (Yale University) laboratories for reagents. Yale Nucleus Club and BMB Club for helpful discussions. This work was supported by: NIH R01 (GM131004) and NSF CAREER (1846010) to S.B. Additional support is by NIH (T32 GM722345) to J.W.C.R, Anderson Postdoctoral Fellowship to S.L., and NIH R35 (GM128666) to M.V.A. and the Alfred P. Sloan Foundation to M.V.A.

## Contributions

J.W. Carrasquillo Rodríguez and S. Bahmanyar conceived the project. J.W. Carrasquillo Rodríguez performed most of the experiments and data analysis. O.U. performed experiments for ER expansion, nuclear solidity and lipin 1 localization in the CTDNEP1-HA stable variants. S. Gao performed experiments for protein purification, size exclusion chromatography and p-NPP analysis of CTDNEP1/NEP1R1 in-vitro with M.V.A. supervision. S. Lee wrote the IJ Macro and Python scripts for LD analysis. J.W. Carrasquillo Rodríguez and S. Bahmanyar wrote the manuscript with input from other authors. S. Bahmanyar supervised the project.

## Materials and Methods

### Mammalian cell lines

U2 OS, Cos 7, DLD-1 and HEK293-T cells were obtained the source specified. Cells were grown at 37°C in 5% CO_2_ in DMEM low glucose (Gibco 11885) supplemented with 10% heat inactivated FBS (F4135) and 1% antibiotic-antimycotic (Gibco 15240112). Cells were cultured without antibiotics during transfections, RNAi, and treatments for experiments. Cells were used for experiments before passage 25. Cells were tested for mycoplasma upon initial thaw and generation of new cell lines (Southern Biotech 13100-01), and untreated cells were continuously profiled for contamination by assessment of extranuclear DAPI/Hoechst 33258 staining.

### Stable cell line generation

U2OS CTDNEP1^KO^ + CTDNEP1 stable cell lines (CTDNEP1(V233E)-HA, Nem1HR- CTDNEP1-HA, CTDNEP1ΔAH-HA, CTDNEP1-EGFP, CTDNEP1(V233E)-EGFP) were generated by retroviral transduction, and bulk populations of cells were used for experiments. Retroviruses were generated by transfecting HEK293T cells with pCG-gag-pol, pCG-VSVG and pMRX-CTDNEP1 using Lipofectamine 2000. The retroviruses were recovered 48hrs post-transfection, filtered using a 0.22um PVDF syringe filter and used to transduce U2OS CTDNEP1^KO^ cells. After 48hrs of infection, cells were placed under 7.5 µg/mL blasticidin or 300µg/mL G418 selection for ∼1 week or until control cells where dead, then frozen and/or used for experiments. Cells were continuously cultured in 7.5 µg/mL blasticidin.

### Transfection and RNAi

Transfections of cells for fix or live imaging were performed with Lipofectamine 2000 (Thermo Fisher Scientific 11668) in Opti-MEM (Gibco 31985) using a 1:2 ratio of DNA:lipofectamine with DNA concentrations ranging from 0.05-0.3 μg DNA per cm^2^ of growth surface. Briefly, DNA and lipofectamine were added to 10 μl OptiMEM per cm^2^ of growth surface in separate borosilicate glass tubes (Thermo Fisher Scientific STT-13100-S). After 5 minutes incubation, DNA solution was added to lipofectamine solution. After 15 minutes, DNA:lipofectamine mix was added dropwise to cells plated 16-24 hrs prior to transfection in fresh antibiotic-free media (1 ml/9.6 cm^2^ growth surface). Media was exchanged for antibiotic-free media after 6 hours.

Transfections of cells for immunoblotting were performed with Polyjet (SL100688) in high glucose DMEM (11965092) using a 1:3 ratio of DNA:Polyjet with DNA concentrations ranging from 0.1 μg DNA per cm^2^ of growth surface. Briefly, DNA and Polyjet were added to 10 μl high glucose DMEM per cm^2^ of growth surface in separate borosilicate glass tubes (Thermo Fisher Scientific STT-13100-S). After 5 minutes incubation, Polyjet solution was added to DNA solution. After 10 minutes, DNA:Polyjet mix was added dropwise to cells plated 16-24 hrs prior to transfection in fresh antibiotic-free media (1 ml/9.6 cm^2^ growth surface). Media was exchanged for antibiotic-free media after 6 hours.

For experiments involving transient CTDNEP1 and/or NEP1R1 overexpression, pcDNA3.0 was used as an empty vector negative control.

RNAi was performed using Dharmafect 1 (Horizon Discovery T-2001) in Opti-MEM according to the manufacturer’s protocol for 20nM. RNAi performed for immunoblot or imaging was analyzed after 48-72hrs.

### Plasmid generation

#### General note

Insertion of gene sequence was conducted either by using restriction enzymes from New England Biolabs, Gibson Assembly (New England Biolabs E5510S) or In-Fusion HD Cloning Plus (Takara 638909). Site-directed mutagenesis was performed by QuikChange Site-directed Mutagenesis Kit (Agilent 200523). Multi site-directed mutagenesis was performed by QuikChange Multi site-directed Mutagenesis Kit (Agilent 200513). Successful cloning was confirmed by sequencing for all constructs.

### Point mutagenesis

CTDNEP1 point mutagenesis was modified from pcDNA or pMRX CTDNEP1-HA or CTDNEP1-GFP using Quickchange Mutagenesis to make the following mutations: V233E, M220A/V233A. NEP1R1 point mutagenesis was modified from pRK5-Flag-NEP1R1 using Quickchange Mutagenesis to make the following mutations: F30E, F30A. For CTDNEP1 region deletions, PCR was performed removing the region of interest followed by In-Fusion (Takara 638909).

### Region deletions and chimeras

For deletion of CTDNEP1’s N-terminus, PCR was performed using primers to amplify the plasmid without the N-terminus, followed by In-Fusion. For Nem1HR-CTDNEP1, PCR was performed with primers deleting the AH and overhangs coding for *yeast* Nem1 hydrophobic region, followed by T4 ligation (M0202).

### Stable cell cloning

pMRX-EGFP-Blast was enzyme digested with BamHI and NotI-HF and gel purified to remove EGFP. pcDNA-CTDNEP1-EGFP were PCR amplified and T4 ligated into the opened backbone of pMRX-Blast to make pMRX-CTDNEP1-EGFP)-Blast. pMRX-CTDNEP1-Blast was subsequently cloned for making point mutations or region deletions has described previously.

### Treatments

Drug/compound treatment was done as follows: Cycloheximide 100 µg/mL; MG132 30µM; CB- 5083 5µM; Oleic acid 200µM; Torin 1 250nM; Bafilomycin A1 100µM. Vehicle control was prepared by diluting the same amount of DMSO as the drug treatment counterpart.

### Cycloheximide treatment assay

Cell media was exchanged with 100 µg/mL cycloheximide. After the indicated time, cells were collected for western blot. Using Image lab, band intensity was measured using the volume tool. Mean intensity was subtracted from background and immunoblot intensity of HA-tag was quantified and normalized to that of α-tubulin, and is shown as % of the value relative to 0 hr.

### Quantitative real-time PCR

RNA was harvested using the RNeasy Mini kit (Qiagen 74104) using the manufacturer’s protocol, using Qiashredder columns (Qiagen 79654) for tissue homogenization and with additional RNase-free DNase (Qiagen 79254) treatment after the first RW1 wash and subsequently adding another RW1 wash. RNA was eluted with RNAse-free water and diluted to 50 ng/μl. RNA was subject to reverse transcription using the iScript Reverse Transcription Supermix (Bio-Rad 1708840) with 400 ng RNA per reaction. The subsequent cDNA was diluted 1:5 for RT-qPCR. cDNA was analyzed for RT-qPCR using the iTaq universal SYBR Green Supermix (Bio-Rad 1725120). Cycle threshold values were analyzed using the Δ ΔCt method. Statistical testing was performed on ΔCt values.

### Immunofluorescence

Cells were washed 2x with warm PBS and fixed in 4% paraformaldehyde (+0.1% glutaraldehyde for ER structure analyses) in PBS for 15 min, permeabilized in 0.5% Triton X-100 for 5 min (or 30mg/mL Digitonin for 10min in ice), then washed 3 times with PBS and blocked in 2% BSA in PBS for 30 min. Samples were transferred to a humidity chamber and incubated with primary antibodies in 2% BSA in PBS for 1 hour at room temperature with rocking. Samples were washed with PBS 3 times for 5 min, then incubated with secondary antibodies in 2% BSA in PBS for 1 hour at room temperature in the dark with rocking. Samples were then washed with PBS 3 times for 5 min in the dark. Coverslips were mounted with ProLong Gold Antifade reagent + DAPI (Thermo Fisher P36935) and sealed with clear nail polish. For samples treated with goat primary antibodies, 2% normal donkey serum (Sigma D9663) was used in place of 2% BSA.

### Immunoblot

Lysis buffers used: 0.1% Triton X-100, 50 mM NaF, 1mM EDTA, 1 mM EGTA, 10 mM Na2HPO4,50 mM b-glycerophosphate, 1 tablet/50 ml cOmplete protease inhibitor cocktail, pH 7.4, RIPA buffer (25 mM Tris pH 7.4, 1% NP-40, 0.5% sodium deoxycholate, 0.1% SDS, 150 mM NaCl, and 1 tablet/50 ml cOmplete Mini protease inhibitor cocktail (Roche 11836153001)). Cell lysates were removed from growth surfaces by scraping with a rubber policeman after incubation in lysis buffer or by adding lysis buffer to cell pellets collected by trypsinization and centrifugation at 300xg for 5 min followed by 1-2 PBS washes. Lysates were homogenized by pushing through a 23G needle 30 times and then centrifuged at >20,000xg for 10 min at 4C, then protein concentration was determined using the Pierce BCA Protein assay kit (Thermo Scientific 23225). 20-30 mg of whole cell lysates/lane were run on 8-15% polyacrylamide gels dependent on target size, and protein was wet transferred to 0.22 mm nitrocellulose. Ponceau S staining was used to visualize transfer efficiency, then washed with TBS or DI water; then, membranes were blocked in 5% nonfat dry milk or BSA in TBS for 1 hour. Membranes were then incubated with primary antibodies in 5% milk or BSA for 1-2 hours at room temperature or overnight at 4C with rocking. Membranes were washed 3 times for 5 min in TBS-T, then incubated with anti-HRP secondary antibodies in 5% milk or BSA in TBS-T for 1 hour at room temperature with rocking. Membranes were washed 3 times for 5 min in TBS-T. Clarity or Clarity Max ECL reagent (Bio-Rad 1705060S, 1705062S) was used to visualize chemiluminescence, and images were taken with a Bio-Rad ChemiDoc or ChemiDoc XRS+ system. Exposure times of images used for analysis or presentation were maximum exposure before saturation of pixels around or within target bands.

Antibody concentrations used: Mouse anti-a tubulin DM1A 1:5000; Mouse anti-Flag 1:4000; Rabbit anti-HA 1:1000; Rabbit anti-lipin 1 1:2000; mouse anti-myc 1:1000; rabbit anti-Sun2 1:1000; mouse anti-GFP 1:1000; mouse anti-GAPDH 1:1000; rabbit anti-LC3B 1:1000; rabbit anti-NRF2 1:1000. all secondaries 1:10000.

### Immunoprecipitation

Cell where pelleted by trypsinization and centrifugation at 300x g for 5 min followed by 1x PBS wash, after pelleting the cells, they were lysed by adding lysis buffer (0.1% Triton X-100, 50 mM NaF, 1mM EDTA, 1 mM EGTA, 10 mM Na_2_HPO_4_, 50 mM β-glycerophosphate, 1 tablet/50 ml cOmplete protease inhibitor cocktail (Roche 11836153001), pH 7.4). Lysates were homogenized by pushing through a 23G needle 30 times and then centrifuged at >20,000x g for 10 min at 4°C. Preconjugated anti-HA magnetic beads (Thermo Scientific 88836) or pre-conjugated anti-Flag magnetic beads (Sigma M8823) were washed twice with TBST and equilibrated in lysis buffer without detergent. 10% of the total volume of lysed cells was transferred to a new tube labelled input, the remaining 90% volume was added to equilibrated beads and incubated for 2 hr rocking at 4°C. Anti-HA beads were then washed twice with TBST and 4x loading dye was added to denature the beads and load samples to SDS-PAGE gel. Anti-Flag beads were washed twice with TBS-T, incubated for 30min with a Flag peptide (ApexBio A6001) and later boiled/loaded into SDS-PAGE gel.

### pNPP assay

100mM pNPP was prepared in 50mM HEPES, pH6.7, 100mM NaCl, 10mM MgCl2, 10mM beta-mercaptoethanol. The reactions were carried out in 96-well plate by mixing 95μL pNPP solution with 5μL enzyme (0.1μM final concentration), and the absorbance at 405nm were monitored with SpectraMax M2e Microplate Readers (Molecular Devices) at 30sec interval at ambient temperature for 30min.

### In vitro interaction of His6-MBP-CTDNEP1ΔAH with NEP1R1

Purified His6-MBP-CTDNEP1ΔAH or His6-MBP-CTDNEP1ΔAH(V233E) (12uM, final concentration) were mixed with NEP1R1 or NEP1R1(F30E) (53uM, final concentration), incubated on ice for 3h, the mixture was centrifuged at 15,000rpm for 15min at 4 °C. The supernatant was loaded onto Superdex200 increase (10/300). Phosphatase activity of each fraction were analyzed by pNPP assay.

### Live cell imaging

For live imaging, cells were plated in ibidi 2 well imaging chambers (ibidi 80287) or ibidi 8 well imaging chambers (ibidi 80827) with DIC lid (ibidi 80055). Samples were imaged in a CO_2_-, temperature-, and humidity-controlled Tokai Hit Stage Top Incubator. Objectives were also heated to 37°C. For CO_2_-controlled imaging, the imaging media used was Fluorobrite DMEM (Gibco A1896701) supplemented with 10% FBS.

### Microscopy

Samples for fix imaging of endoplasmic reticulum, digitonin vs triton permeabilization and samples for live-cell microscopy of localization of CTDNEP1 with other markers were imaged on an inverted Nikon Ti microscope equipped with a Yokogawa CSU-X1 confocal scanner unit with solid state 100-mW 488-nm and 50-mW 561-nm lasers, using a 60×1.4 NA plan Apo objective lens (or 10x 0.25 NA ADL objective with 1.5x magnification), and a Hamamatsu ORCA R-2 Digital CCD Camera.

Samples for live cell imaging of CTDNEP1-EGFP stable siRNA or CTDNEP1 for cells treated with oleate for function in lipid droplet analysis were imaged on an inverted Nikon Ti Eclipse microscope equipped with a Yokogawa CSU-W1 confocal scanner unit with solid state 100 mW 405, 488, 514, 594, 561, 594, and 640 nm lasers, using a 60x 1.4 NA plan Apo objective lens and/or 20x plan Fluor 0.75 NA multi-immersion objective lens, and a prime BSI sCMOS camera.

### Oleate treatment and lipid droplet staining

Oleic acid (oleate) was conjugated to fatty acid free BSA at a molar ratio of 6:1. For experiments of CTDNEP1 localizing to lipid droplets, 200μM oleate and 0.5 μM BODIPY 558/568 C12 was added in Fluorobrite DMEM (Gibco A1896701) supplemented with 10% FBS and imaged after 24hrs. For functional experiments of lipid droplets number and area, cells were treated with 200μM oleate and 1 μM BODIPY 558/568 C12 for 24hrs, next day media was changed to Fluorobrite DMEM supplemented with 10% FBS and imaged immediately.

### Protein modeling

Protein complex structure prediction was performed using ChimeraX plug-in into google collab for complex structure prediction. Protein prediction outputs where modelled/analyzed using PyMOL education software.

### Amphipathic helix predictions

HeliQuest was used to generate a helical wheel projection and to obtain hydrophobic moment <µH> and net charge Z values, which yielded a discriminant factor D = 0.944 (<µH>) + 0.33 (z) as described in HeliQuest (https://heliquest.ipmc.cnrs.fr/HelpProcedure.htm).

### Conservation score analysis

For CTDNEP1 amphipathic helix alignment amino acid sequences were obtained from uniprot and species were chosen such that they include *S. cerevisiae (P38757), C. elegans (Q20432), D. melanogaster (Q9VRG7), H. sapiens (O95476), M. musculus (Q3TP92), D. rerio (Q5U3T3)*. For NEP1R1 sequence alignment species were chosen to include *S. cerevisiae (P18410), S. pombe (Q9USQ0), C. elegans (Q9XXN3), D. melanogaster (Q8T0B1), H. sapiens (Q8N9A8), M. musculus (Q3UJ81), D. rerio (Q561X0)*. A multiple sequence alignment and phylogenetic tree were generated using Clustal Omega with the default settings.

### Image analysis

Image analysis was performed using FIJI/ImageJ. For all scoring phenotypes quantified by categorization were scored blindly. Images were blinded for analysis using the ImageJ

Macro ImageJ Filename_Randomizer, cells where randomized and analysis was done blindly. For scoring CTDNEP1 involvement in lipid droplet biogenesis, cells where automatically threshold and segmented using a Python macro, followed by manual color thresholding and utilization of the Analyze Particles tool in FIJI/ImageJ with 0.9 - 1.0 particle circularity has the only restriction for LD number.

ER phenotypes were additionally quantified with percent abundance of cytoplasmic calnexin signal. For cells with the entire ER captured within 0.3-0.5 mm interval z stacks, 8-bit maximum intensity projections were made of the whole field of view. To ensure the different ER morphologies were all accounted for after thresholding, the 8-bit max projections were subject to unsharp masking with a radius of 2 and mask of 0.6. The max intensity projection was thresholded using the Huang threshold of object fuzziness (Huang and Wang, 1995). The cell border and nuclear border for each cell were manually traced using ER fluorescent signal. For percent occupancy of the cytoplasm by ER membranes, the percent of pixels within the nucleus-free cell area that were calnexin (+) was measured.

### Statistical analysis

GraphPad Prism 8 was used for all statistical analysis. Continuous data was tested for normality using a Shapiro-Wilk test. In imaging experiments where phenotypes of individual cells are scored, n refers to individual cells. All N refer to experimental repeats. p values, Fisher’s exact tests for ER expansion phenotype and One-way Anova with Dunnet’s multiple comparisons test for lipid droplet phenotype.

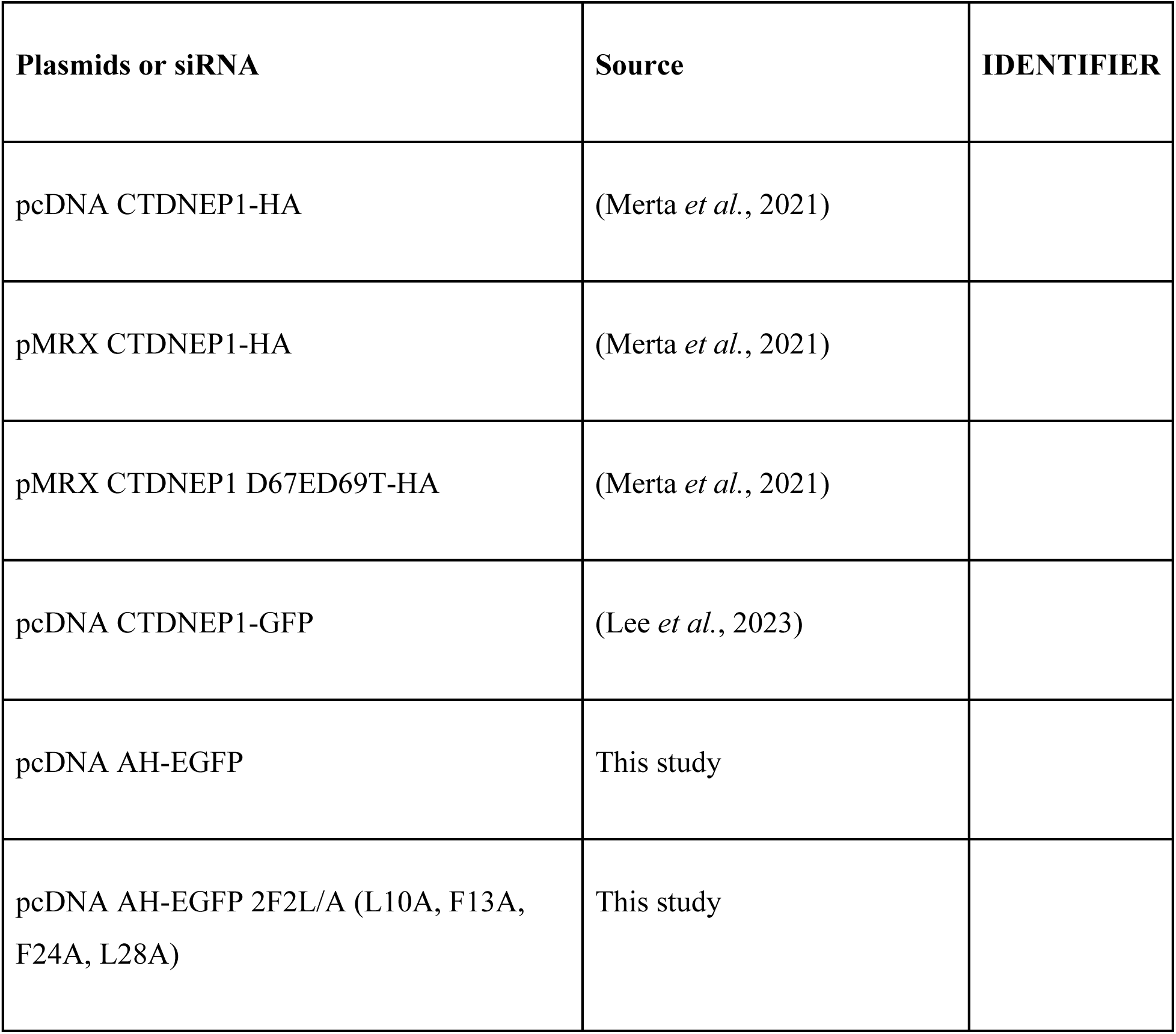
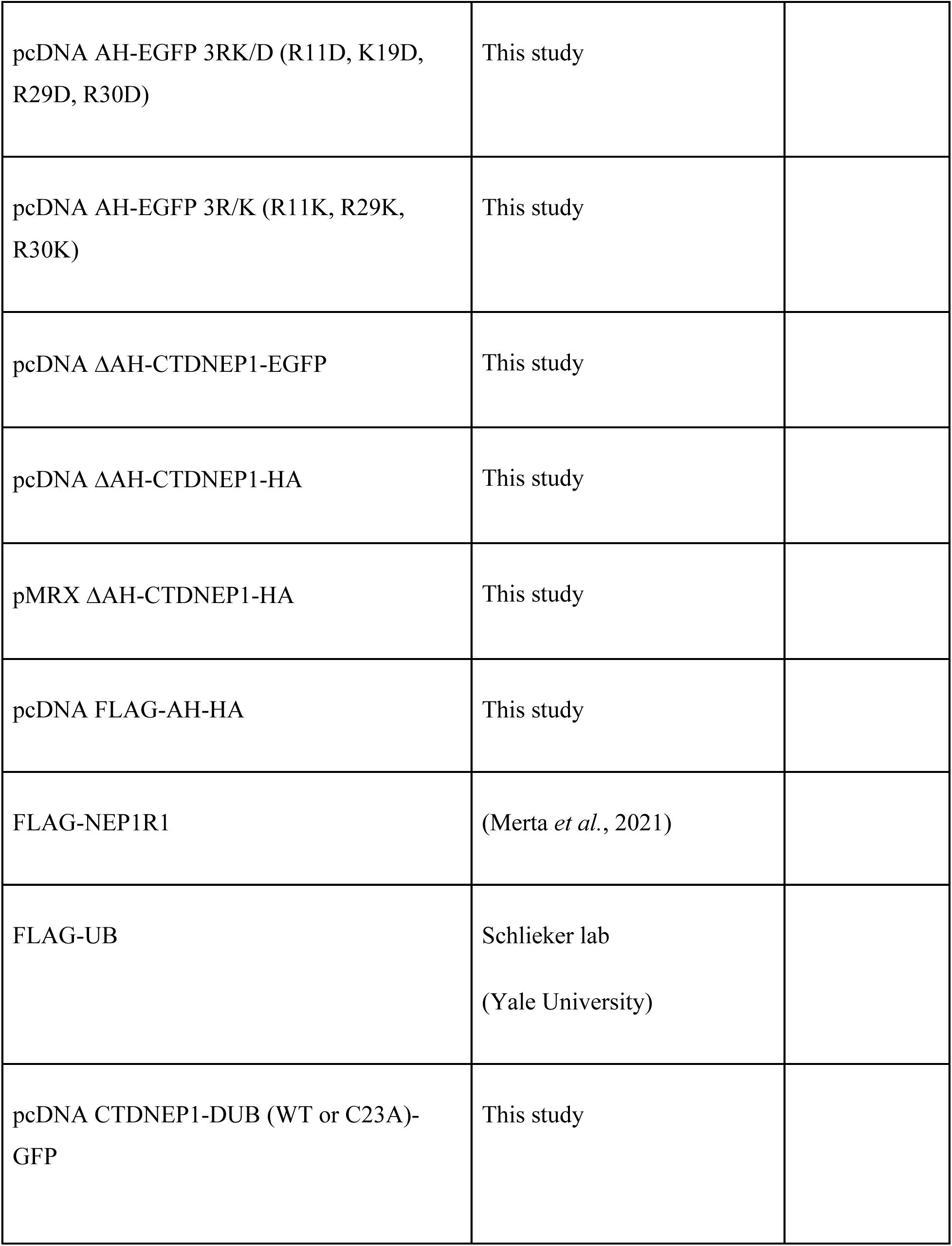
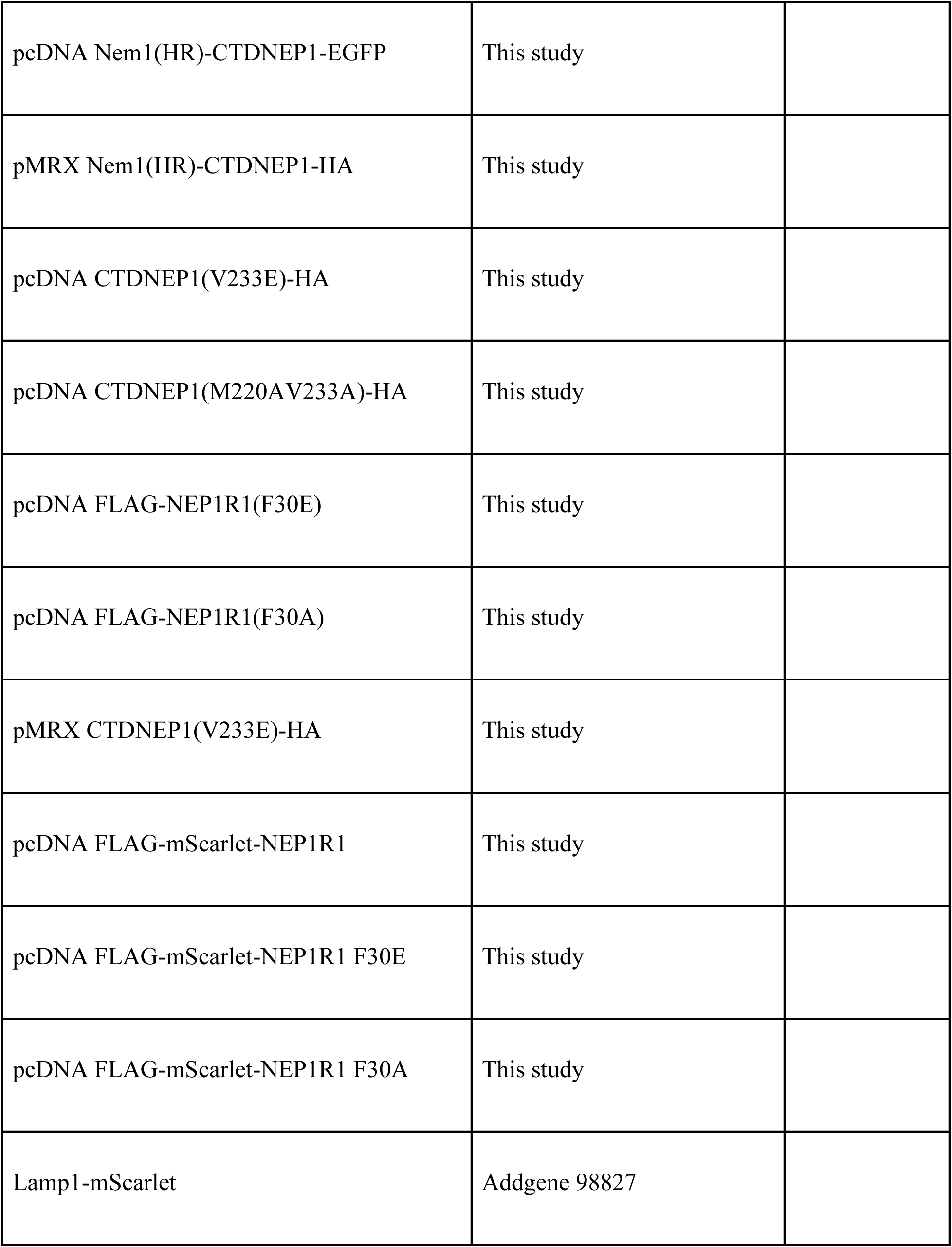
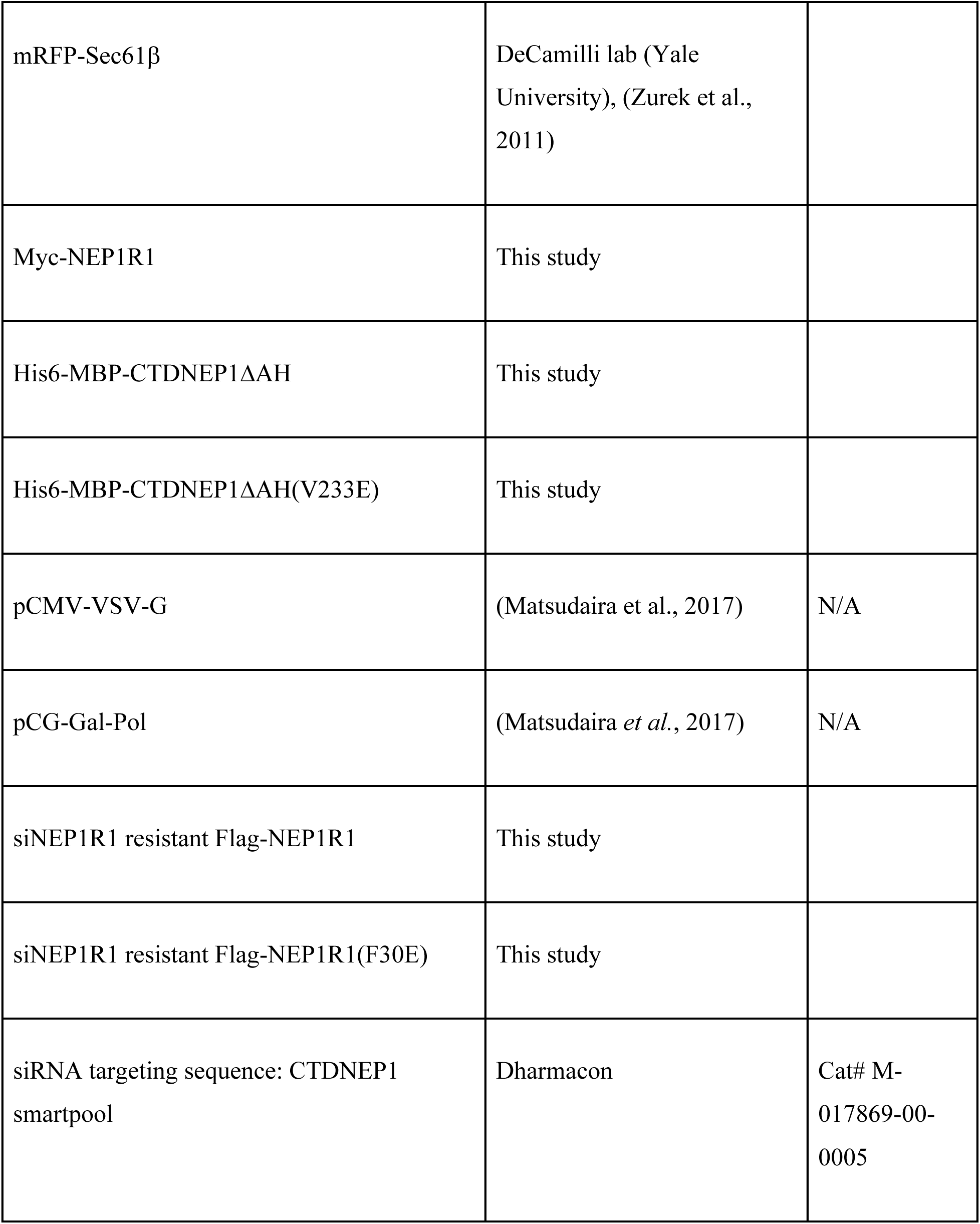
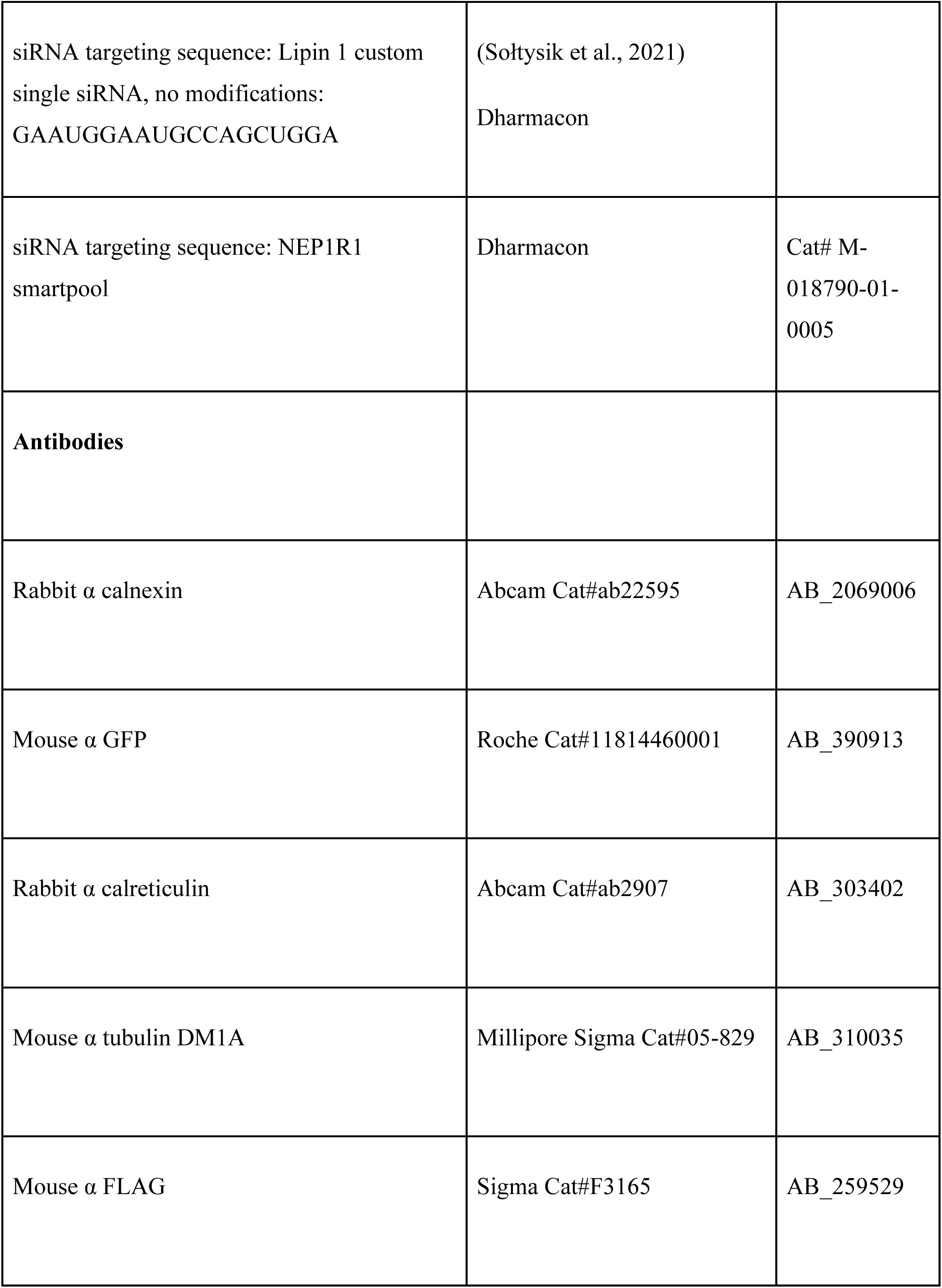
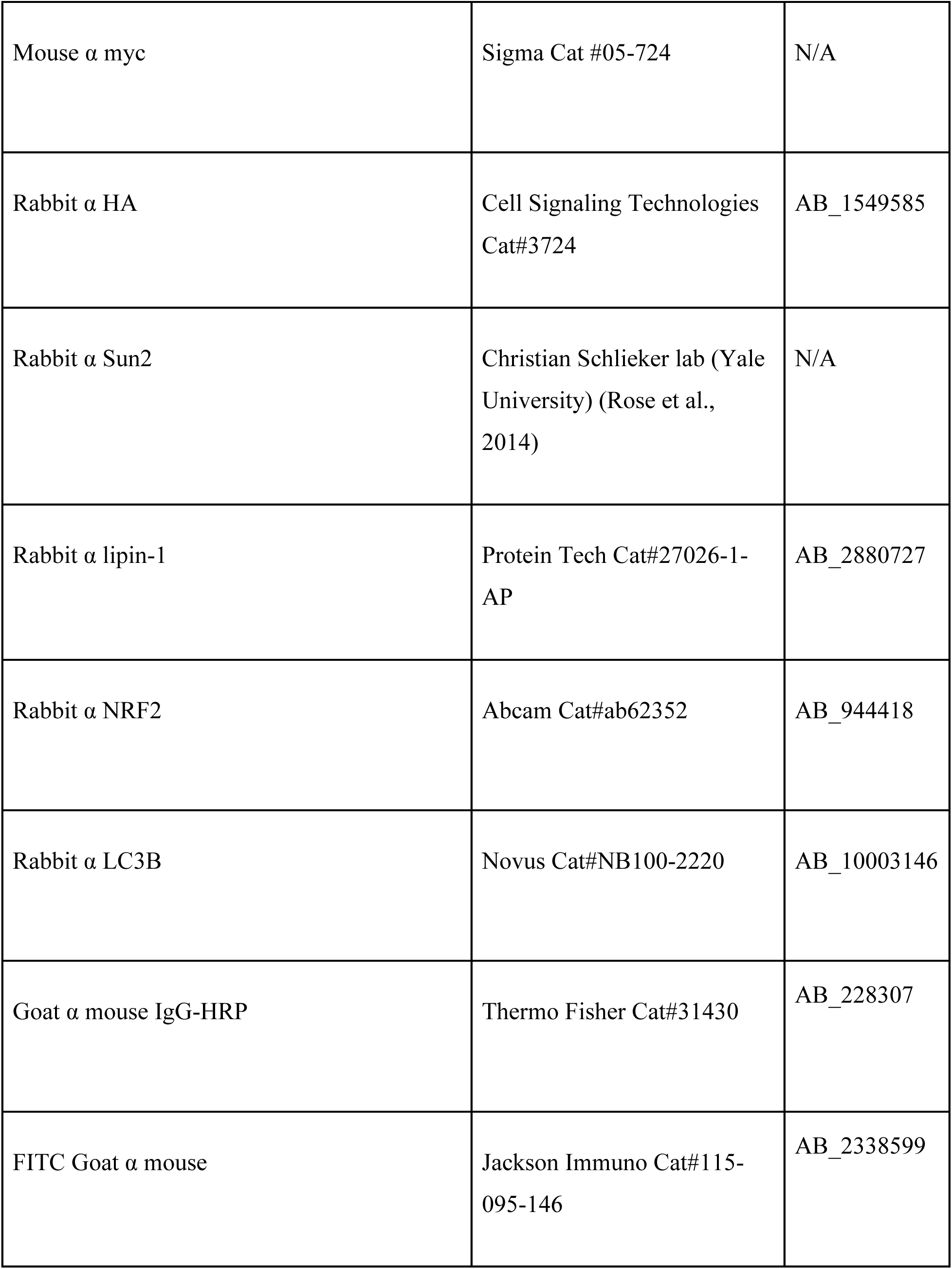
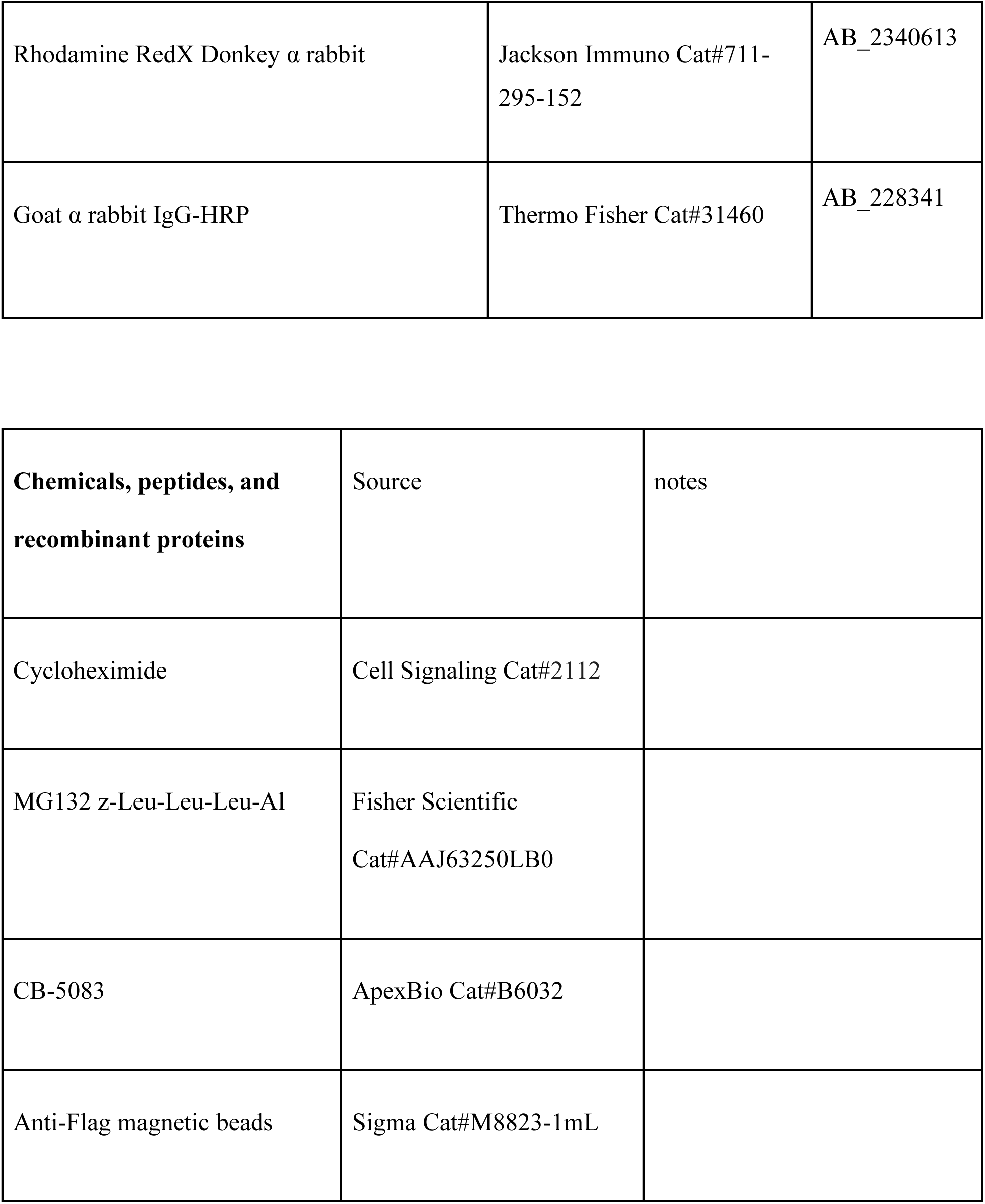
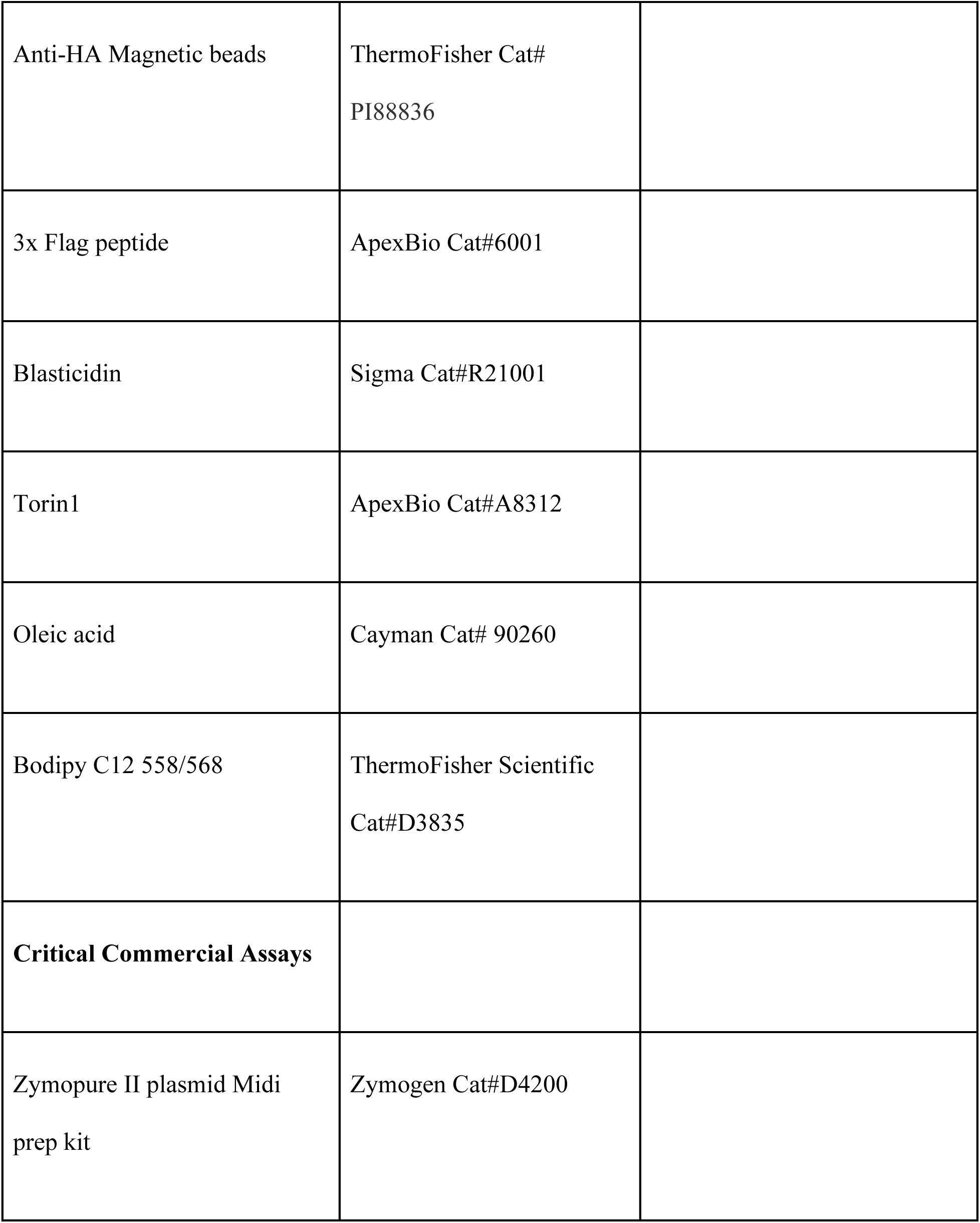
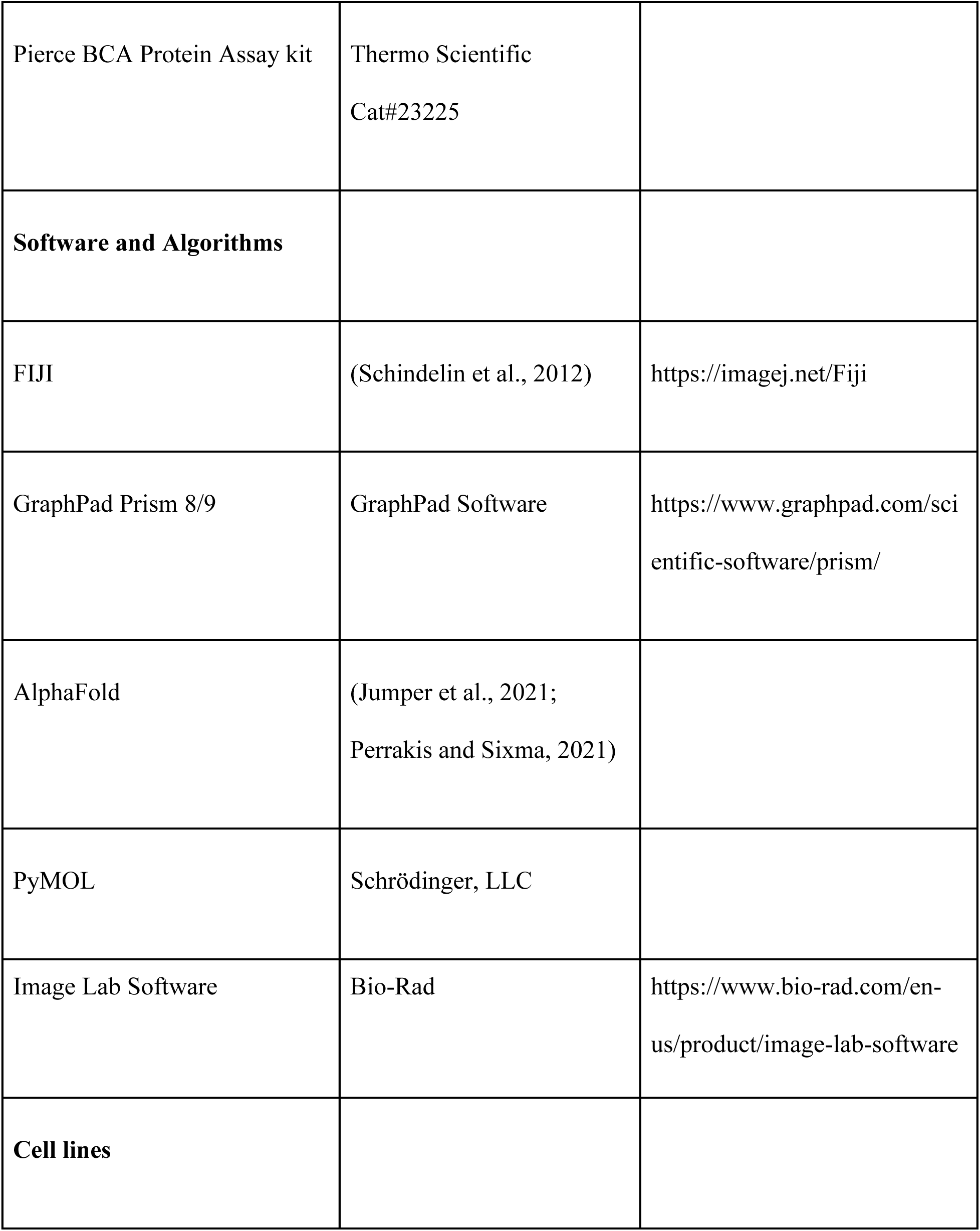
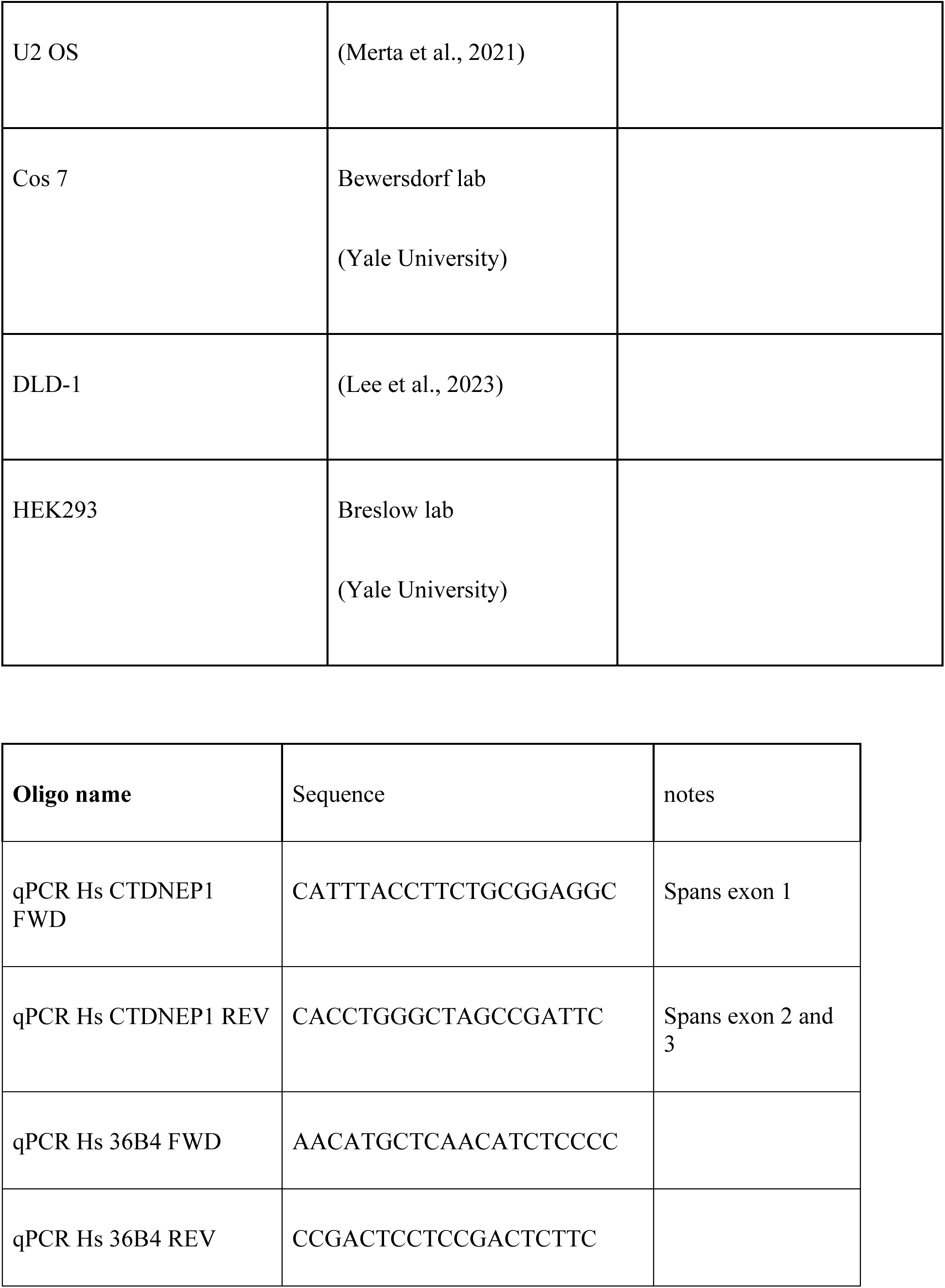
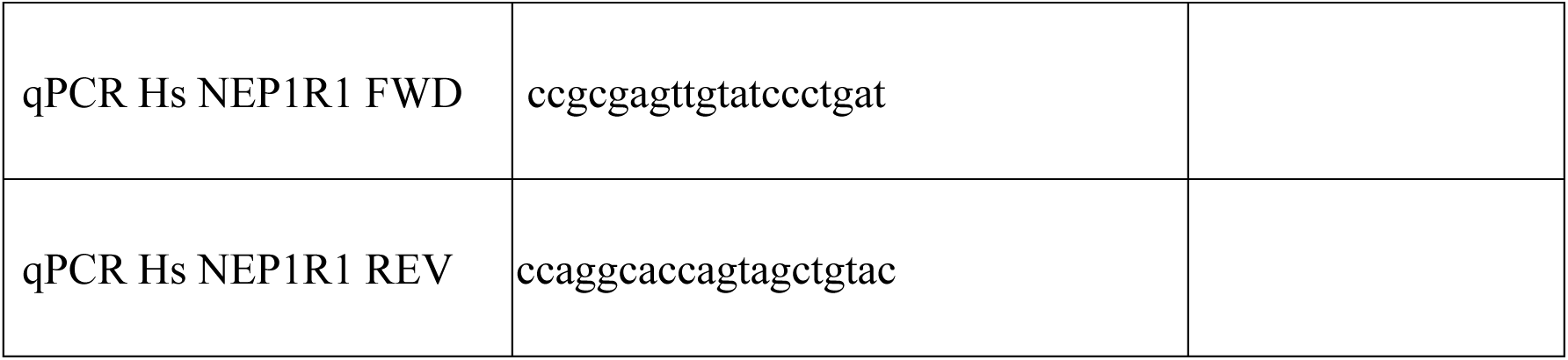

---

**Supplemental Figure 1, support for main Figure 1.**
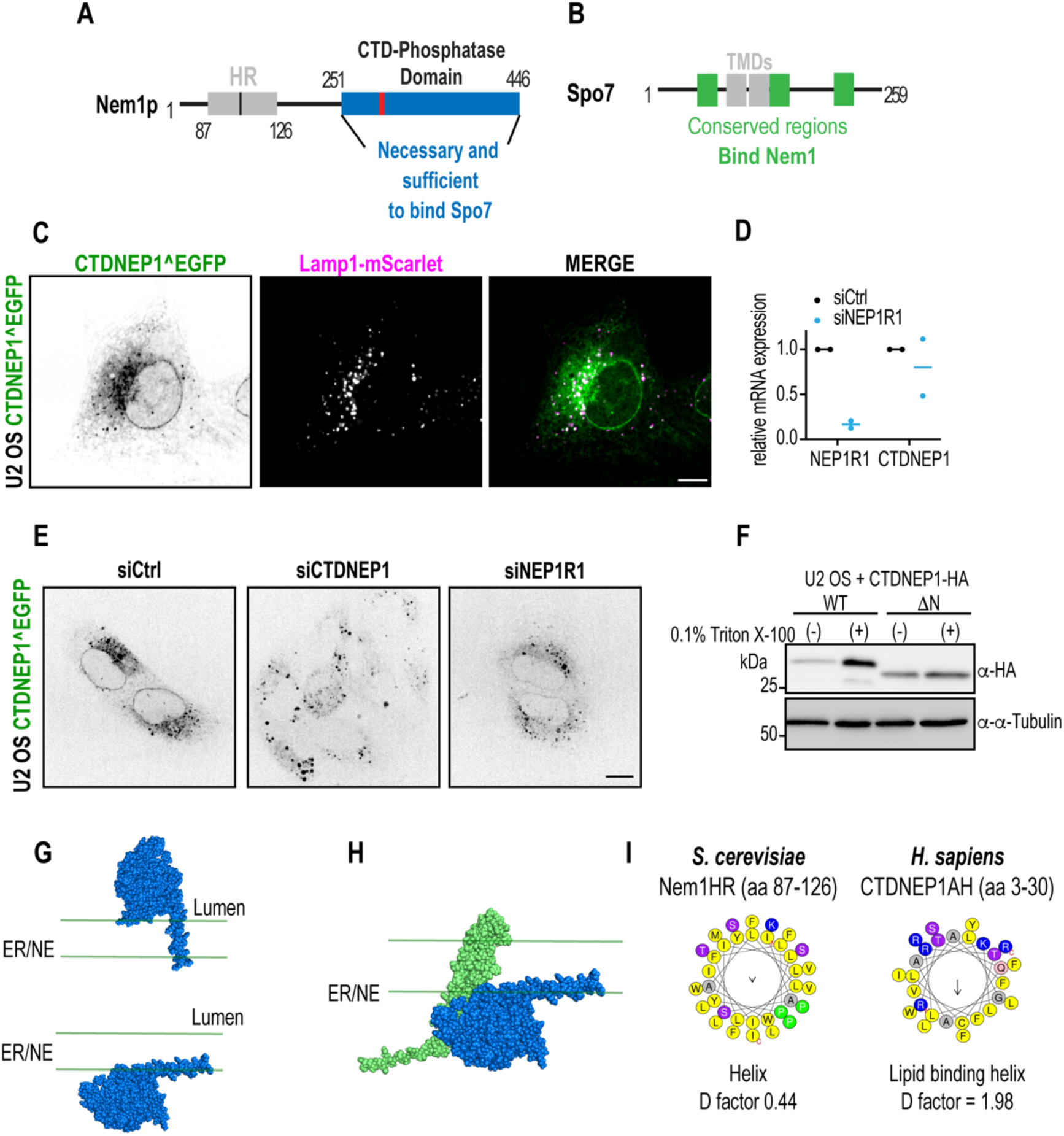
A-B) Schematic representation of A) Nem1 and B) Spo7 protein domain. C) Representative spinning disk confocal images of CTDNEP1^EGFP and Lamp1-mScarlet in living cells. D) qRT-PCR of CTDNEP1^EGFP cells for genes indicated, shown as fold change of expression relative to mean of control values. E) Representative spinning disk confocal images of CTDNEP1^EGFP in living cells RNAi depleted of the indicated genes. F) Immunoblot of U2 OS whole cell lysates transfected with CTDNEP1-HA variants and extracted with phosphatase buffer ± 0.1% Triton X-100. G) Schematic of CTDNEP1 orientation within the membrane directed by its N-terminus. H) AlphaFold protein complex prediction of CTDNEP1 (blue) and NEP1R1 (green) oriented at membranes by the transmembrane domains of NEP1R1. I) Schematic of helical wheel projection of N-terminus of human CTDNEP1 (aa 3-30) compared to hydrophobic region of Nem1 (aa 87-126) generated by HeliQuest with the discrimination factor (D factor). For all: scale bars, 10µm, N = 2 independent experiments.

**Supplemental Figure 2, support for main Figure 2.**
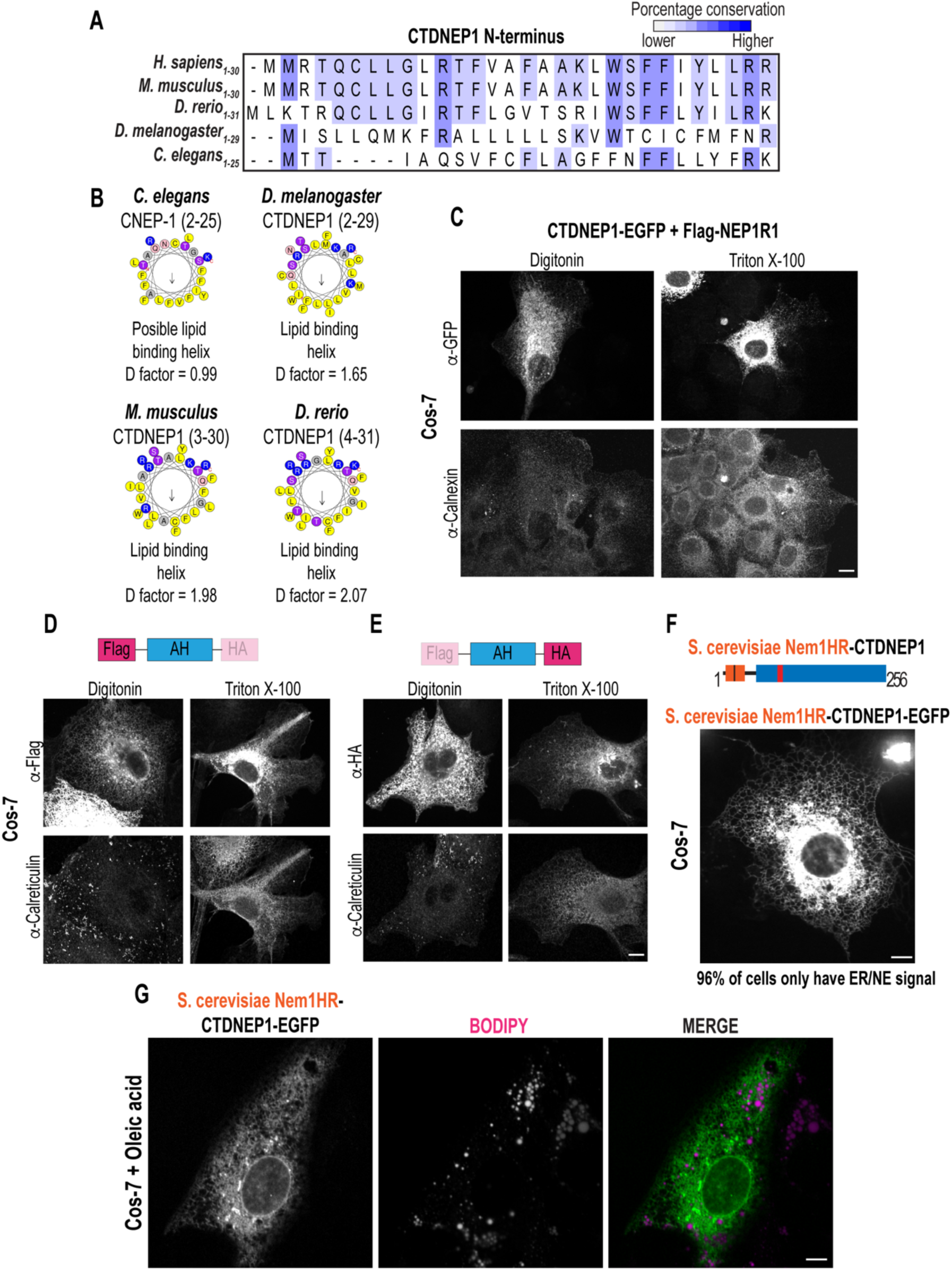
A) Multiple sequence alignments of the N-terminus of CTDNEP1 in metazoans. B) Schematic of helical wheel projection of N-terminus of *C elegans, D. melanogaster, M. musculus and D. rerio* CTDNEP1 generated by HeliQuest with the discrimination factor (D factor). C) Representative spinning disk confocal max projection images of GFP and calreticulin immunostained from Cos-7 cells transfected with CTDNEP1- EGFP and Flag-NEP1R1, under the indicated permeabilization conditions. D-E) Representative spinning disk confocal images of calreticulin and HA or Flag immunostained from Cos-7 cells transfected with Flag-AH-HA, under the indicated permeabilization conditions. F) Top: Schematic representation of CTDNEP1 chimera with N-terminus (aa 3-30) replaced by Nem1 hydrophobic region (87-126), Bottom: Representative spinning disk confocal image of Nem1HR-CTDNEP1-EGFP in living Cos-7 cells. N = 2 independent experiments G) Representative spinning disk confocal images of Nem1HR-CTDNEP1-EGFP in living Cos-7 cells treated with 200µM Oleic acid and stained with Bodipy C12 for 24hrs. N = 2 independent experiments. For all: scale bars, 10µm.

**Supplemental Figure 3, support for main Figure 3.**
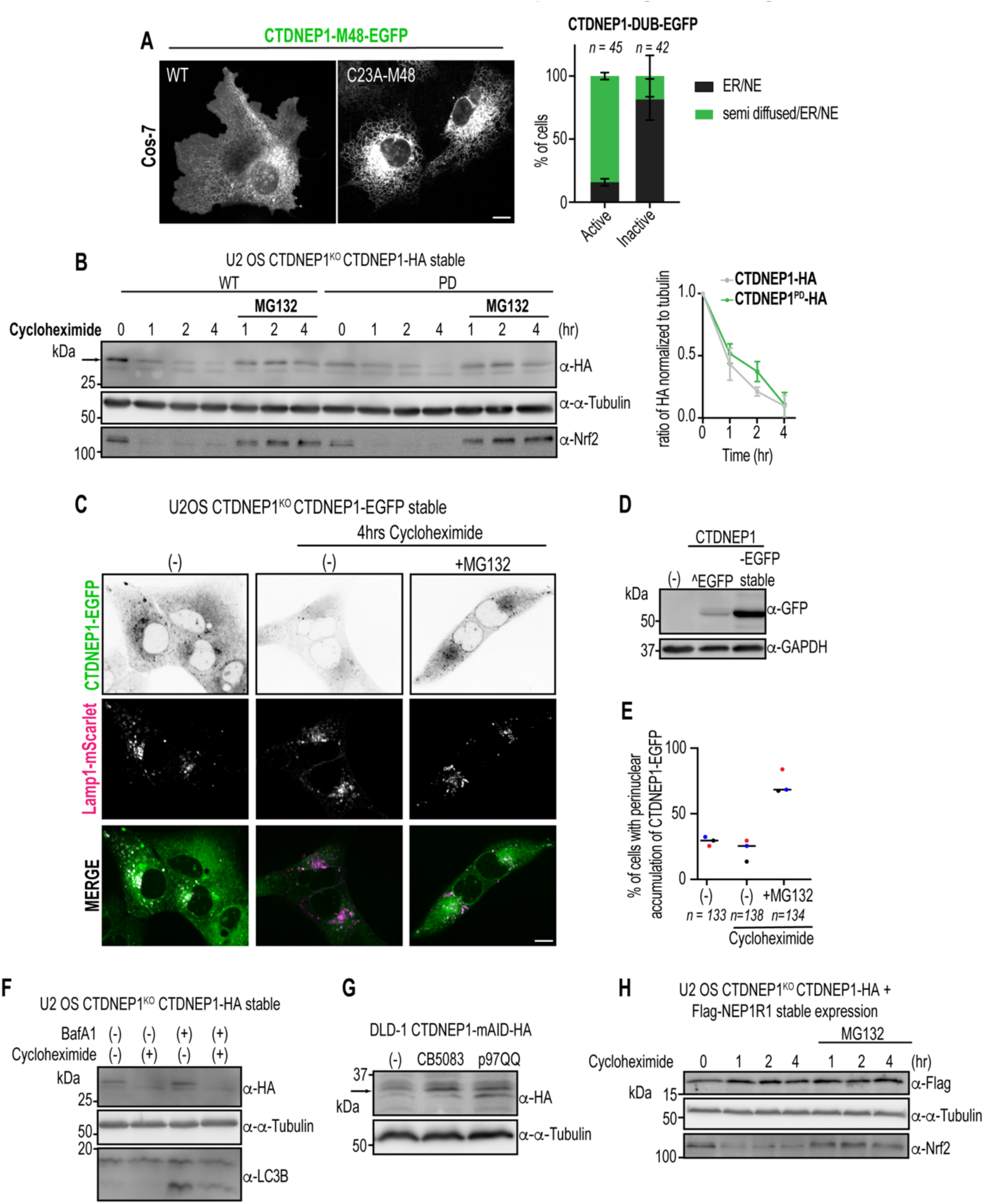
A) Left: Representative spinning disk confocal images of CTDNEP1-DUB-EGFP in living Cos-7 cells. Right: Plot: blind quantification of CTDNEP1–DUB-EGFP localization. Mean ± SD shown, n = individual cells counted, N = 3 independent experiments. B) Left: Immunoblot of whole cell lysates of CTDNEP1 deleted cells stably expressing CTDNEP1-HA variants treated with cycloheximide ± MG132 for the indicated time. Right: Plot, represents mean intensity value of top band of CTDNEP1-HA relative to its initial timepoint (0hr) normalized to tubulin. Mean ± SD shown. N = 3 independent experiments. C) Representative spinning disk confocal images of CTDNEP1- EGFP in living cells under the indicated conditions. D) Immunoblot of whole cell lysates of U2 OS CTDNEP1^EGFP or CTDNEP1 deleted cells stably expressing CTDNEP1-EGFP. N = 2 independent experiments. E) Plot: blind quantification of CTDNEP1-EGFP perinuclear accumulation from Figure S3C. N = 3 independent experiments and mean shown. n = individual cells counted. F) Immunoblot of whole cell lysates of CTDNEP1 deleted cells stably expressing CTDNEP1-HA treated with bafilomycinA1 and/or cycloheximide for 4 hours. N = 2 independent experiments G) Immunoblot of whole cell lysates of DLD-1 CTDNEP1 endogenously tagged with mAID-HA cells, treated with CB-5083 or transfected with p97 QQ. N = 2 independent experiments H) Immunoblot of whole cell lysates of CTDNEP1 deleted cells stably expressing CTDNEP1-HA and Flag-NEP1R1 treated with cycloheximide ± MG132 for the indicated time. N = 2 independent experiments. For all: scale bars, 10µm

**Supplemental Figure 4, support for main Figures 4 and 5.**
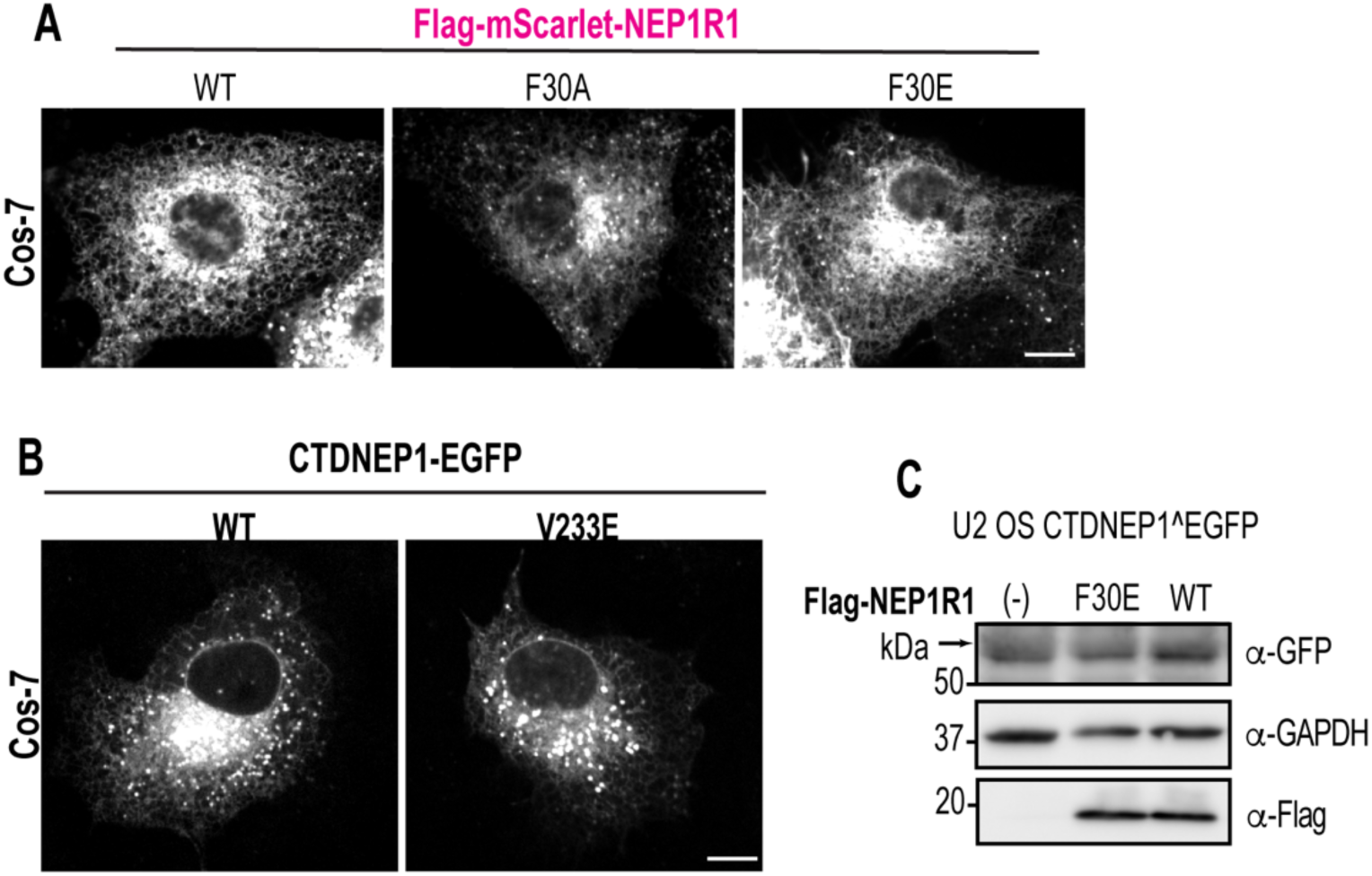
A) Representative spinning disk confocal images of Flag-mScarlet-NEP1R1 variants in transfected Cos-7 living cells. B) Representative spinning disk confocal images of CTDNEP1-EGFP variants in transfected Cos-7 living cells. C) Immunoblot of whole cell lysates of CTDNEP1^EGFP cells transfected with Flag-NEP1R1 variants. For all: scale bars 10µm, N = 2 independent experiments.

**Supplemental Figure 5, support for main Figure 6.**
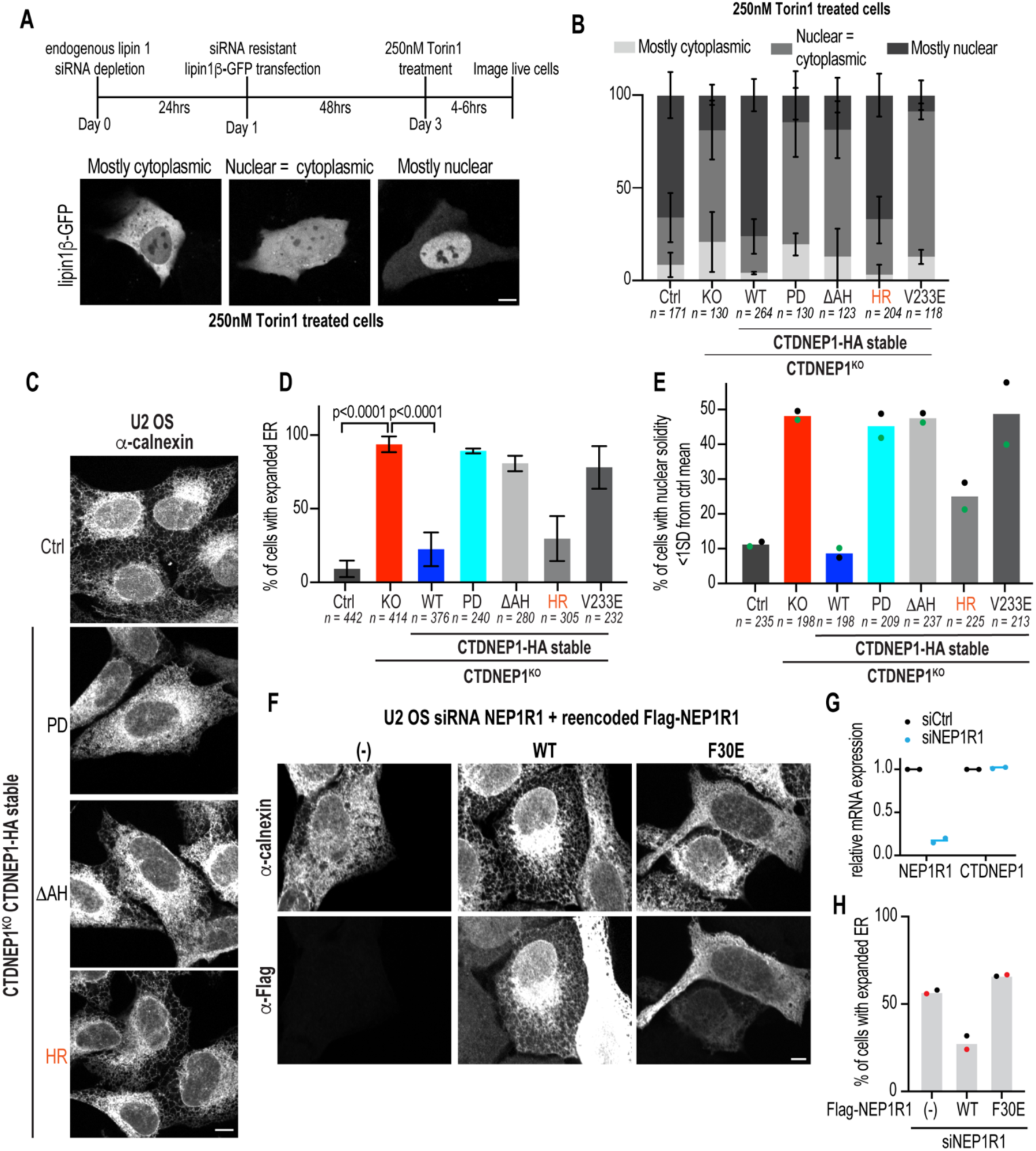
A) Top: Schematic of experimental design for lipin1β-GFP localization, Bottom: Representative spinning disk confocal images of lipin1β- GFP in living cells. B) Plot: blind quantification of lipin1β-GFP localization in the different CTDNEP1 backgrounds treated with 250nM Torin1 for 4-6 hours. Means ± SD shown, N = 3 independent experiments. C) Representative spinning disk confocal images of calnexin staining in fixed cells. D) Plot: blind quantification of percentage of cells with expanded ER in indicated cells. Mean ± SD shown. N = 3 independent experiments, p value, Fisher’s exact tests. E) Plot: automated quantification of incidence of nuclei with solidity value <1SD from control mean solidity. Mean and N = 2 independent experiments shown. F) Representative spinning disk confocal images of calnexin and Flag immunostaining in fixed cells. G) qRT-PCR of U2 OS cells RNAi depleted of NEP1R1 for the genes indicated, shown as fold change of expression relative to mean of control values. H) Plot: blind quantification of percentage of cells with expanded ER. Mean and N = 2 independent experiments shown. For all: scale bars, 10µm, n = individual cells counted.

